# Interaction of GAT1 with sodium ions: from efficient recruitment to stabilisation of substrate and conformation

**DOI:** 10.1101/2023.10.10.561652

**Authors:** Erika Lazzarin, Ralph Gradisch, Sophie M.C. Skopec, Leticia Alves da Silva, Chiara Sebastianelli-Schoditsch, Dániel Szöllősi, Julian Maier, Sonja Sucic, Marko Roblek, Baruch I. Kanner, Harald H. Sitte, Thomas Stockner

## Abstract

The human GABA transporter (GAT1) is a membrane transporter that mediates the reuptake of the neurotransmitter GABA from the synaptic cleft into neurons and glial cells. Dysregulation of the transport cycle has been associated with epilepsy and neuropsychiatric disorders, highlighting the crucial role of the transporter in maintaining homeostasis of brain GABA levels. GAT1 is a secondary active transporter that couples the movement of substrate to the simultaneous transport of sodium and chloride ions along their electrochemical gradients. Using MD simulations, we identified a novel sodium recruiting site at the entrance to the outer vestibule, which attracts positively charged ions and increases the local sodium concentration, thereby indirectly increasing sodium affinity. Mutations of negatively charged residues at the recruiting site slowed the binding kinetics, while experimental data revealed a change in sodium dependency of GABA uptake and a reduction of sodium affinity. Simulation showed that sodium displays a higher affinity for the sodium binding site NA2, which plays a role in stabilisation of the outward-open conformation. We directly show that the presence of a sodium ion bound to NA2 increases the stability of the closed inner gate and restrains motions of TM5. We find that sodium is only weakly bound to NA1 in the absence of GABA, while the presence of the substrate strengthens the interaction due to the completed ion coordinating shell, explaining cooperativity between GABA and sodium.

## Introduction

γ-aminobutyric acid (GABA) is the main inhibitory neurotransmitter in the central nervous system (CNS) (Krnjević and Schwartz, 1967). Release of GABA from the presynaptic neuron into the synaptic cleft triggers hyperpolarization of the postsynaptic neuron through activation of GABA_A_ receptors (Ghit et al., 2021). GABA is removed from the synaptic cleft by several GABA transporters, namely GAT1, GAT2, GAT3 and the betaine/GABA transporter BGT1. These are sodium- and chloride-dependent secondary active transporters from the SLC6 family (Bhatt et al., 2023; Kanner, 1978; Kristensen et al., 2011) with GAT1 being the first cloned member of the family (Guastella et al., 1990). Within the GABA transporter family, the presynaptic GAT1 is the main neuronal GABA uptake transporter. Dysregulation of GAT1 function has been associated with epilepsy and neuropsychiatric disorders (Fischer et al., 2022; Kasture et al., 2023), highlighting the crucial role of GAT1 in maintaining neurotransmitter homeostasis in the brain (Bhat et al., 2021; Schousboe et al., 2004; Schousboe and Madsen, 2017)

GAT1 consists of 12 transmembrane helices (TM) (Motiwala et al., 2022) and shares the pseudo-symmetric inverted repeat motif (Forrest et al., 2008) of the “LeuT fold” with the other SLC6 family members, including the fold defining bacterial small amino acid transporter LeuT (Yamashita et al., 2005), the Drosophila dopamine transporter dDAT (Penmatsa et al., 2013), and the human serotonin transporter hSERT (Coleman et al., 2016), which was the first structure for a human transporter of the SLC6 family. TM3, 4, 8 and 9 constitute the core of the rigid scaffold domain, while TM1, 2, 6 and 7 assemble into the more mobile bundle domain. The motion of the bundle domain relative to the scaffold domain allows for alternating access to the central substrate binding site S1 (Forrest et al., 2008).

Very recently, several inward-facing GAT1 structures and a substrate, sodium and chloride-bound inward-occluded structure were published (Motiwala et al., 2022; Nayak et al., 2023). These conformations together with an AlphaFold2 model of the outward-facing state have allowed to formulate the translocation cycle for GAT1 (Nayak et al., 2023; Zhu et al., 2023) which consists of outward-open, occluded and inward-open states. The conformations of the outward-open and the occluded structures are very similar to the respective conformations observed in other SLC6 family members (Coleman et al., 2016; Singh et al., 2008; Yamashita et al., 2005; Yang and Gouaux, 2021), confirming shared structural changes across the SLC6 family. In contrast, SLC6 transporter structures differ in the conformation of TM1a of the inward-facing state, whereby for GAT1 several conformations of TM1a were observed (Motiwala et al., 2022; Nayak et al., 2023; Zhu et al., 2023). For LeuT we have shown that the large motions of TM1a as observed in the micelle solubilized crystal structure (Krishnamurthy and Gouaux, 2012) are incompatible within the membrane environment (Sohail et al., 2016), suggesting that the conformation of TM1a is especially sensitive to the environment.

The concentrative uptake of GABA is enabled by the co-transport with two sodium ions and one chloride (Bhatt et al., 2023; Hilgemann and Lu, 1999; Keynan and Kanner, 1988). The chloride binding site and the two sodium binding sites (NA1 and NA2) are located in the centre of the transporter, next to the unwound regions on TM1 and TM6, which form the substrate binding site (S1) together with TM3 and TM8 (Nayak et al., 2023; Zhu et al., 2023; Zomot et al., 2007). Structural considerations on LeuT (Krishnamurthy and Gouaux, 2012; Yamashita et al., 2005), MD simulations (Szöllősi and Stockner, 2022) and biochemical studies (Tavoulari et al., 2016) indicated that the sodium bound to the NA2 (Na2) is involved in stabilisation of the outward-open state by connecting TM1 of the bundle with TM8 of the scaffold. The sodium ion bound to the NA1 (Na1) is exclusively interacting with the bundle domain. It stabilises the bound substrate by directly interacting with its carboxyl group in GAT1 and LeuT.

Here we present direct evidence for the mechanism by which the two sodium ions exert their function in GAT1. Using a set of 60 simulations starting from the sodium-free outward-open GAT1 state, we investigated sodium binding to and the response of GAT1 to the presence of sodium ions. We observed comparable probability for initial binding of sodium to either NA1 or NA2. Importantly, only Na2 was stably bound, while we observed several unbinding events of Na1 from NA1 and in some cases a transfer of the sodium to NA2. Analysis of the simulations identified a clear sodium entry path with a sodium recruiting site at the entry to the extracellular vestibule and revealed a temporary site in the S1. The role of the recruitment site that includes two negatively charged residues (D281 and E283) on TM6a is to attract sodium ions, which increases the local sodium concentration and accelerates sodium binding. We find that Na1 stabilises GABA in the S1 by including its carboxyl moiety into the first interaction shell of Na1, holding the carboxyl moiety in the same position in space as the carboxylate of the conserved aspartate (D98 in SERT, D79 in DAT) of the monoamine transporters. Na2 is directly linked to the mobility of the bundle domain as Na2 binding reduces its mobility as well as increases the stability of the inner gate between TM1, TM5 and TM6.

## Results

### The path taken by sodium to reach NA1 and NA2

Sodium ions reach the ion binding sites NA1 and NA2 through the outer vestibule. Binding to the outward-open conformation of SLC6 transporters is a fast process in the nano to microsecond time range (Hilgemann and Lu, 1999; Bicho and Grewer, 2005; Hasenhuetl, Freissmuth and Sandtner, 2016; Szöllősi and Stockner, 2021), therefore allowing for an unbiased direct simulation approach. Using 60 independent simulations, we repeatedly observed that sodium ions spontaneously entered the outward-open vestibule of GAT1. To characterise the path that sodium ions take for reaching the S1 and for binding to NA1 and NA2, we derived an averaged sodium density with a 0.1 nm spatial resolution from the sodium positions observed in all simulations. These sodium density maps (Figure 1a) highlight the regions with a high probability of observing sodium ions. The density analysis shows that sodium ions initially accumulate at the rim of the outer vestibule. This sodium recruitment site comprises two negatively charged amino acids (D281 and E283) that are located at the beginning of TM6a (Figure 1b). The high sodium density at the recruitment site shows that sodium can more likely be found at the recruitment site compared to the extracellular solution. This locally higher concentration increases the probability of binding, because sodium ions can leave the recruitment site in two directions: either (i) enter the open outer vestibule towards the S1, attracted by the negative field in the S1 and the outer vestibule, or (ii) detach from GAT1 and diffuse into the external solution.

**Figure 1:**
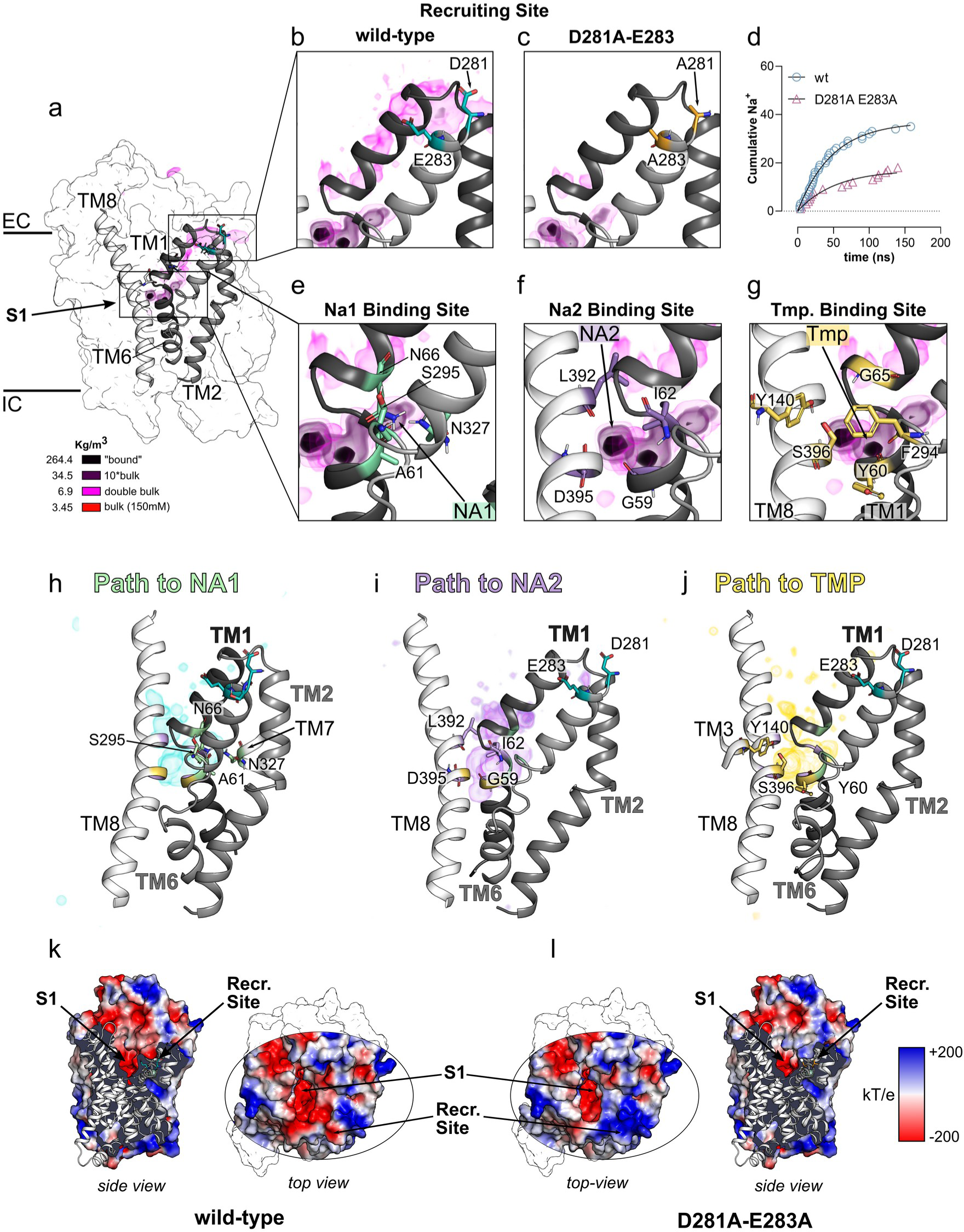
Sodium binding and GAT1 electrostatics. **a)** The GAT1 surface is shown in white, with transmembrane helices TM1, TM2, and TM6 highlighted in grey. Averaged density of sodium is shown in close proximity to GAT1 in a side view. The sodium density is represented as isovolume, colour-coded according to the legend and shown in three layers above bulk (double bulk, 10* bulk, bound), as indicated by the legend. The trajectories were fitted to the C*α* atoms of the transmembrane domain. The residues D281 and E283 (Recruitment Site) are illustrated as cyan sticks. **b**) The focus zooms in on the recruitment site in GAT1-WT and **c)** the D281A-D283A mutant, showing helices of TM1, TM2, TM6, and TM8 along with the recruitment site residues. Sodium density from the wild-type and the double alanine (D281A and E283A) mutant simulations is colour-coded according to the legend. **d)** Quantification of sodium binding events calculated as cumulative sodium ions entering the vestibule. **e)** Zoom to the S1 showing Ion density and surrounding residues of NA1, **f)** NA2 and **g)** the temporary binding sites. **h)** Separation of ion position observed for sodium ions binding directly to NA1 (green), **i)** directly to NA2 in purple or **j)** first residing in the temporary binding site (yellow) before reaching NA1 or NA2. All residues of the respective binding sites are represented as sticks and coloured accordingly. **k)** Top view and side view of the electrostatic potential of GAT1 wild-type mapped onto its surface. **l)** Top and side view of the electrostatic potential of the double alanine mutant (D281A and E283A).

The salt bridge of the extracellular gate between R69 and D451 was defined as the structural threshold for considering a sodium ion within the vestibule, because once beyond this gate, ions would typically not leave the vestibule, but reach the S1 and bind to NA1 or NA2 (Figure 1e-g). We independently analysed trajectories in which sodium would bind to the NA1 or NA2 and found that the routes sodium ions take through the outer vestibule are indistinguishable. Once in the S1, sodium ions immediately bound to NA1 or NA2, or resided for some time in a temporary pre-binding site within the S1 and only in a second step reached NA1 or NA2 (Figure 1h-j). In some cases, Na1 unbound from the NA1 returning to this temporary site, and occasionally bound to NA2 in a second step. We did not observe unbinding from the NA2. This temporary pre-binding site (Figure 1g,j) for sodium overlaps with the region where the positively charged amino group of the substrate GABA resides in the ligand-bound conformation (Zhu et al., 2023). A similar temporary site and comparable binding kinetics were observed in simulations of direct sodium binding to SERT (Szöllősi and Stockner, 2021). It also overlaps with the region where the positively charged amino group of serotonin was found in SERT (Gradisch et al., 2022; Yang and Gouaux, 2021) suggesting that the temporary binding site is a common feature of SLC6 transporters.

### Role of the sodium recruitment site

Sodium ions typically interact first with the recruiting site at the beginning of the outer vestibule before passing through the outer vestibule and binding to NA1 or NA2. Although negatively charged residues are present in other regions of the extracellular vestibule, and alternative pathways for sodium entry are available, sodium preferentially follows the same path, presumably, because attracted to the recruitment site. Electrostatic potentials (Figure 1h,i) showed the expected strong negative potential in the S1 and the vestibule, but also revealed a strong negative potential at the recruitment site. The residues D281 and E283 are in the centre of this area of negative electrostatic potential. A sequence alignment of all human SLC6 transporters (Figure S1) showed that the negative charge in position 281 is present in 70% of human SLC6 transporters and is an aspartate in all GABA transporters. Position 283 is polar or charged, but the glutamate amino acid is not conserved among GABA transporters. Their mutation in the paralog transporters DAT and SERT were shown to reduce substrate transport or sodium binding (Chen et al., 2004; Kortagere et al., 2013; Szöllősi and Stockner, 2021). *In silico* neutralisation of these charges by alanine mutations (D281A-E283A) collapsed the local negative electrostatic potential, as it changed sign and became weekly positive (Figure 1j,k), supporting the notion of an electrostatic attraction. To explore the impact of D281 and E283 on sodium binding, the double alanine mutation (D281A-E283A) was introduced in GAT1, and ion binding simulations were repeated from the same 60 systems. Consistent with the inference from the electrostatic potential, the density analysis showed a strongly reduced sodium density at the recruitment site (Figure 1c), confirming that sodium ions are electrostatically attracted and that residues D281 and E283 play a critical role. The overall sodium density in the S1, NA1 and NA2 was also lower due to the slower binding kinetics leading to a lower number of sodium binding events.

To determine the importance of the recruitment site for sodium binding, we quantified the kinetics of sodium entering the S1. Figure 1d shows the time dependent cumulative number of simulations in which a sodium ion enters the S1. The data revealed that the double mutant reduced the binding kinetics as compared to wild-type GAT1, thereby showing that the recruitment site accelerates the rate of sodium binding. Consistent with the density analysis, time-dependent distance plots (Figure S2a,b (wild-type) and Figure S2c (double mutant D281-E283) show that sodium ions typically first interact with the recruitment site in wild-type GAT1 before reaching the S1, while no such initial interaction is seen in the double mutant. To verify this prediction *in vitro*, we mutated D281 and E283 to alanine and stably expressed the single mutants (D281A, E283A) and the double mutant in HEK293 cells to experimentally measure uptake and assessed the transport associated currents. The single mutant E283A was expressed similar to wild-type GAT1 (P=0.5423). However, the D281A mutant and D281A-E283A double mutant exhibited a significant reduction in surface expression by approximately 3-fold and 4-fold, respectively (P < 0.0001) (Figure 2a,b). Plasma membrane localisation was clearly detectable for all variants, as visible in the insert of Figure 2a. We quantified intracellular accumulation and membrane expression of the YFP-tagged GAT1 by measuring the mean fluorescence of intracellular and membrane localised GAT1, respectively (Figure 2b). In contrast, the cytosolic fluorescence was similar for wild-type and all three mutants (Figure 2a,b). To verify functional expression of the single mutants (D281A and E283A) and the D281A-E283A variant of GAT1, we measured dose-dependent GABA uptake (Figure 2c). Consistent with surface expression, uptake of GABA was like wild-type (Vmax: 2335.43 ± 571.14 [confidence interval: 2130 - 2508 pmol/10^6^ cells/3 min]) for the E283A mutant (Vmax: 2486.83 ± 292.62[confidence interval: 2368 - 2587 pmol/10^6^ cells/3 min]), approximately 2-fold lower for the D281A mutant (Vmax: 1486.68 ± 135.74[confidence interval: 1434 - 1537 pmol/10^6^ cells/3 min]) and 4-fold lower in the D281A-E283A double mutant (Vmax: 581.17 ± 150.14[confidence interval: 525 - 621 pmol/10^6^ cells/3 min]) (Figure 2c). Also Km values were similar for wild-type GAT1 (Km: 12.00 ± 4.12 µM [confidence interval: 9.11 - 14.36 µM]) and the E283A mutants (Km: 11.11 ± 1.86 µM [confidence interval: 9.66 - 12.38 µM]), the Km for the D281A mutant was smaller (Km: 7.83 ± 1.00 µM [confidence interval: 6.99 - 8.76 µM]) but the difference was statistically not significant (P=0.0582), while the double mutant (Km: 6.01 ± 2.84 µM [confidence interval: 4.26 - 7.36 µM]) showed a significant twofold lower Km (P=0.0054). This data demonstrates that the transport function of GAT1 was not compromised by the mutations and the differences can mainly be associated with reduced expression.

**Figure 2.**
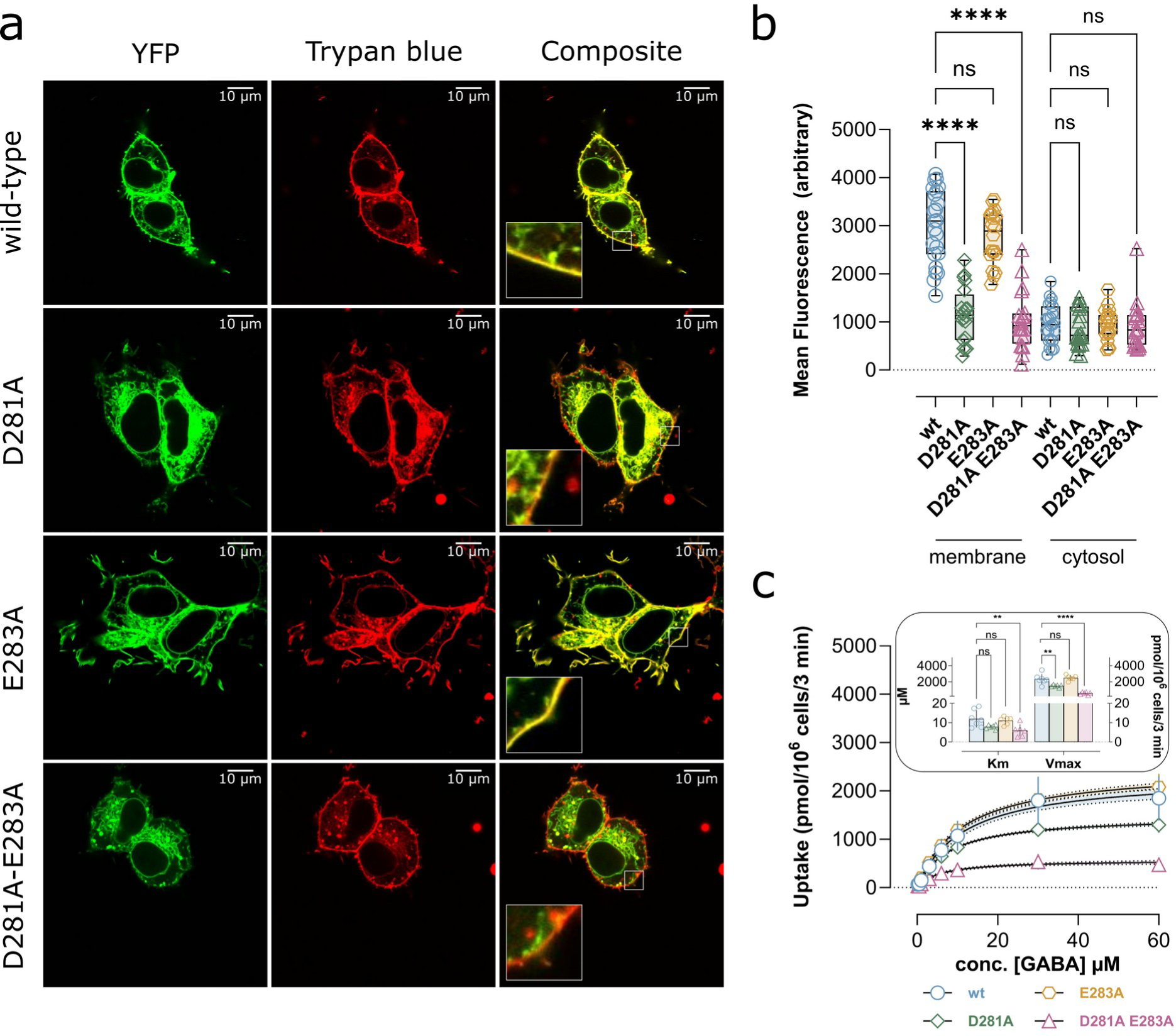
Effect of D281A-E283A mutation. **a)** Representative confocal microscopy images of GAT1 wild-type, single mutants (D281A, E283A) and double alanine mutant (D281A-E283A) stably expressed in HEK293 cells. Transporters were N-terminally tagged with YFP., The cell membrane was stained by trypan blue. Composite images display the superimposition of tagged GAT1 and trypan blue stained membranes. The inserts in the lower left corner show a zoom-in on the cell membrane. **b)** Mean fluorescence of cytosolic and membrane resident YFP-tagged GAT1 of n=23 (wild-type) and n=20 (D281A,E283A,and the double mutant) randomly analysed confocal images per condition. **c)** Concentration-dependent [^3^H]GABA uptake in HEK293 cells stably expressing GAT1 wild-type, single mutants and D281A-E283A. **d)** Insert is depicting the quantification of the Km (µM) and the Vmax (pmol/10^6^ cells/3 min) of the GABA uptake. Data is displayed as mean ± SD. Concentration dependent uptake was measured in triplicates in n=7 (wild-type) and n=6 (D281A, E283A, and D281A-E283A) biologically independent experiments. Single data points for Km and Vmax are the averages for individual experiments. Confidence intervals (95%) for curve fittings are plotted within ± SD and delimited by black dashed lines. After conducting statistics with one-way ANOVA, we performed Dunnett’s post hoc multiple comparison vs. wild-type. Statistical significance was denoted as follows: *=P < 0.05, **=P < 0.01, ***P < 0.001, ****P < 0.0001.

Simulations suggested that the double mutant (D281A-E283A) would affect sodium binding. To unmask a possible effect of sodium on GABA transport, we carried out GABA uptake with increasing concentrations of transport energising sodium. The results (Figure 3a, b) showed that the EC50 of wild-type GAT1 is 46.00 ± 4.51 μM [confidence interval: 42.14 - 49.34 µM] and single charge neutralisation mutants D281A (57.25 ± 2.82 µM [confidence interval: 54.86 - 58.91 µM]) and E283A (52.08 ± 2.06 µM [confidence interval: 50.11 - 54.01 µM]) had a non-significant effect on the sodium dependency of uptake (P=0.129 and P=0.5515, respectively). In contrast, the D281A-E283A double mutant significantly increased the sodium dependency of GABA transport to 80.73 ± 14,28 μM [confidence interval: 70.13 - 84.19 µM] (P < 0.0001). The observed shift of the concentration response curve to the right showed that the Km of sodium dependency of GABA transport is reduced by 43% in double mutant D281A-E283A, verifying the computational predictions that D281 and E283 at the recruitment site effectively increases the local concentration of sodium at the entry to the outer vestibule, which leads to a locally increased sodium concentration and an enhanced sodium binding.

**Figure 3.**
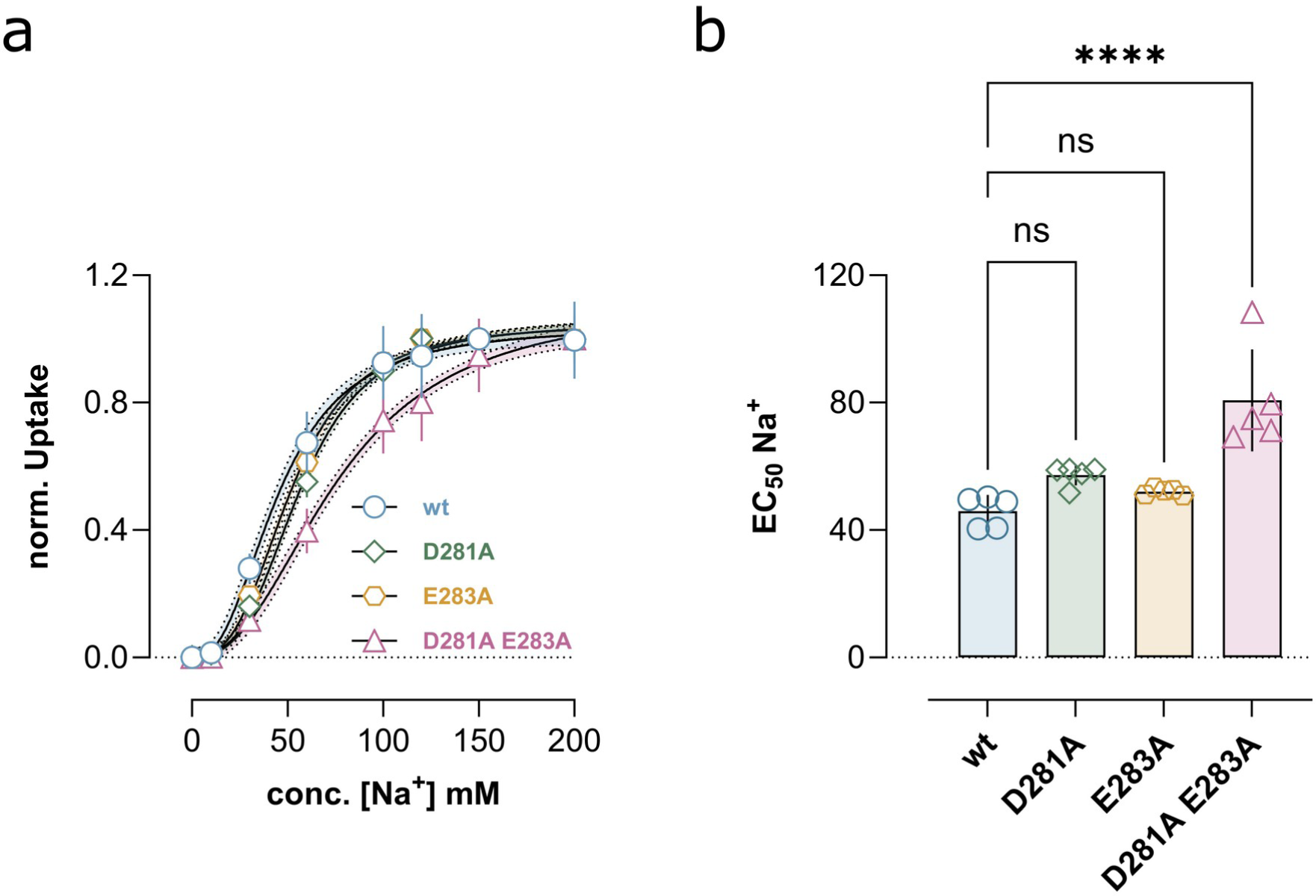
Impact of the charge neutralisation on Na^+^ affinity. **a)** Sodium-dependent [^3^H]GABA uptake curves normalised to maximal uptake. **b)** Sodium EC_50_ values for GAT1 wild-type, D281A, E283A, and the double mutant D281A-E283A observed in panel a. Data is shown as mean ± SD from n=5 biological independent replicas of [^3^H]GABA uptake experiments measured in triplicates. Single EC_50_ data points are the averages for individual triplicates. Confidence intervals (95%) for curve fittings are plotted within ± SD and delimited by black dashed lines. Statistical analysis was performed with one-way ANOVA and Dunnett’s post hoc multiple comparison vs. wild-type. Statistical significance was denoted as follows: *=P < 0.05, **=P < 0.01, ***P < 0.001, ****P < 0.0001.

To functionally characterise the D281A-E283A variant of GAT1, we measured transport function using electrophysiological recordings, as GABA transport by GAT1 is electrogenic. Representative traces of GABA uptake dependent currents are shown for wild-type GAT1 (Figure 4a) and the D281A-E283A double mutant (Figure 4c). The concentration-response of GABA uptake showed for wild-type GAT1 (Figure 4a, b) an EC50 of 18.60 µM [confidence interval: 33.40 µM - 44.23 µM]. The EC50 for the D281A-E283A variant is similar 15.77 µM [confidence interval: 4.87 - 46.97 µM] (Figure 4c, d), confirming that the double mutant affected expression, but not the intrinsic transport function of GAT1 at physiological conditions (Figure 4e).

**Figure 4.**
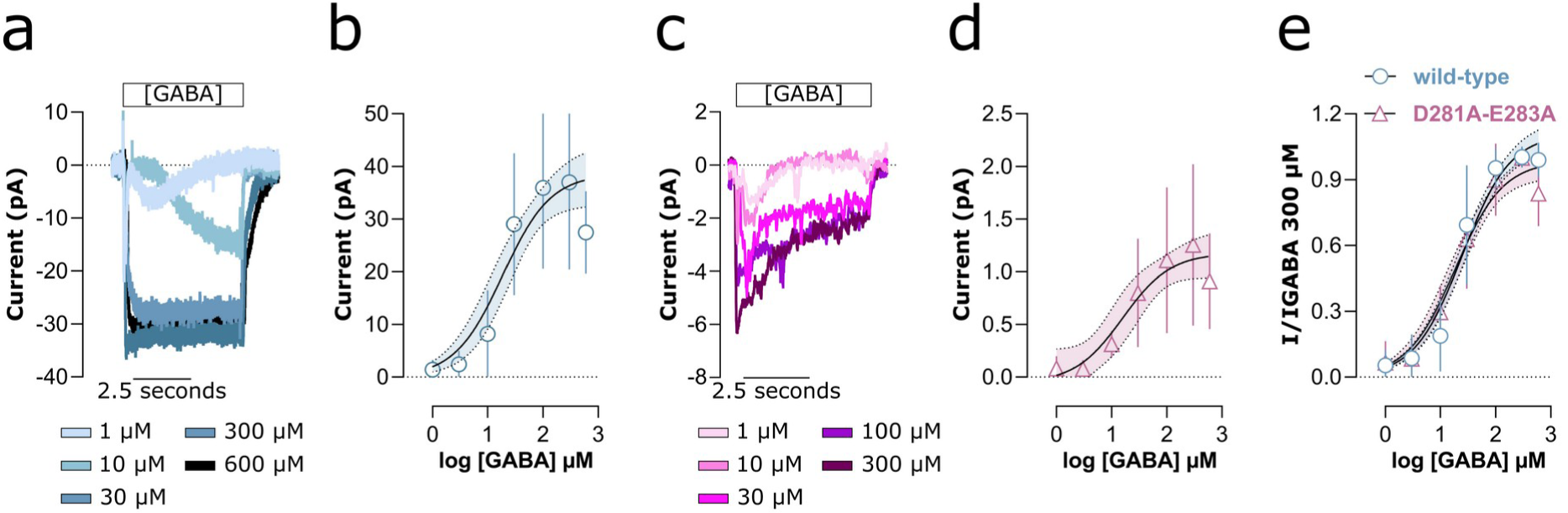
Electrophysiological characterisation of the transport cycle. **a)** Representative traces of whole-cell patch clamp recordings showing GAT1 wild-type mediated currents in response to different GABA concentrations. **b)** Concentration dependent average steady state currents of wild-type GAT1. **c)** Representative traces of whole-cell patch clamp experiments showing concentration-dependent GAT1 D281A-E283A mediated currents (data points are shown as running average (n=11) traces). **d)** Average steady state currents of GAT1 double mutant (D281A-E283A) plotted against the used GABA concentration. **e)** Comparison of concentration dependent average steady state currents of GAT1 wild-type and double mutant, normalised to the respective currents observed at 300 *μ*M GABA. Data are represented as the mean ± SD from n=11 recordings of independent cells per condition. 95% confidence intervals of curve fittings are plotted within ± SD with dashed lines.

### Local structural changes in the sodium binding sites NA1 and NA2

Binding of ligands is frequently accompanied by structural changes in the protein, both in the immediate vicinity of the binding site and at distant locations. Conformational selection and induced fit are two models that are commonly used to describe the structural changes induced by ligand binding. In the conformational selection model (Monod et al., 1965), the protein dynamically interchanges between multiple conformations and the ligand selectively binds and stabilises one conformation. In contrast, the induced-fit model (Koshland JR., 1959) describes the protein to initially exist in a conformation that is not fully complementary to the ligand and it undergoes a ligand-induced conformational change upon binding.

To characterise the structural changes of GAT1 upon sodium binding to NA1 and NA2 (Figure 5a), we first calculated the centre of mass (CoM) of the ion binding site(s), and then measured the average distance between the CoM and all coordinating oxygen atoms (Figure 5b,c, Figure S3) to quantify their compactness. The averaged compactness of NA1 and NA2 was determined in all simulations which showed sodium binding, whereby each simulation was divided into the pre-binding and post-binding part (Figure 5d,e). The mean compactness of NA1 remained unaffected by sodium binding (Figure 5d), while the breadth of the distribution increased slightly, indicating that in GAT1, sodium binding to the NA1 does not induce the same conformational change as observed for SERT (Szöllősi and Stockner, 2021). In contrast, the NA2 (Figure 5e) site showed after sodium binding a clear transition to a more compact geometry, reminiscent of an induced-fit mechanism.

**Figure 5.**
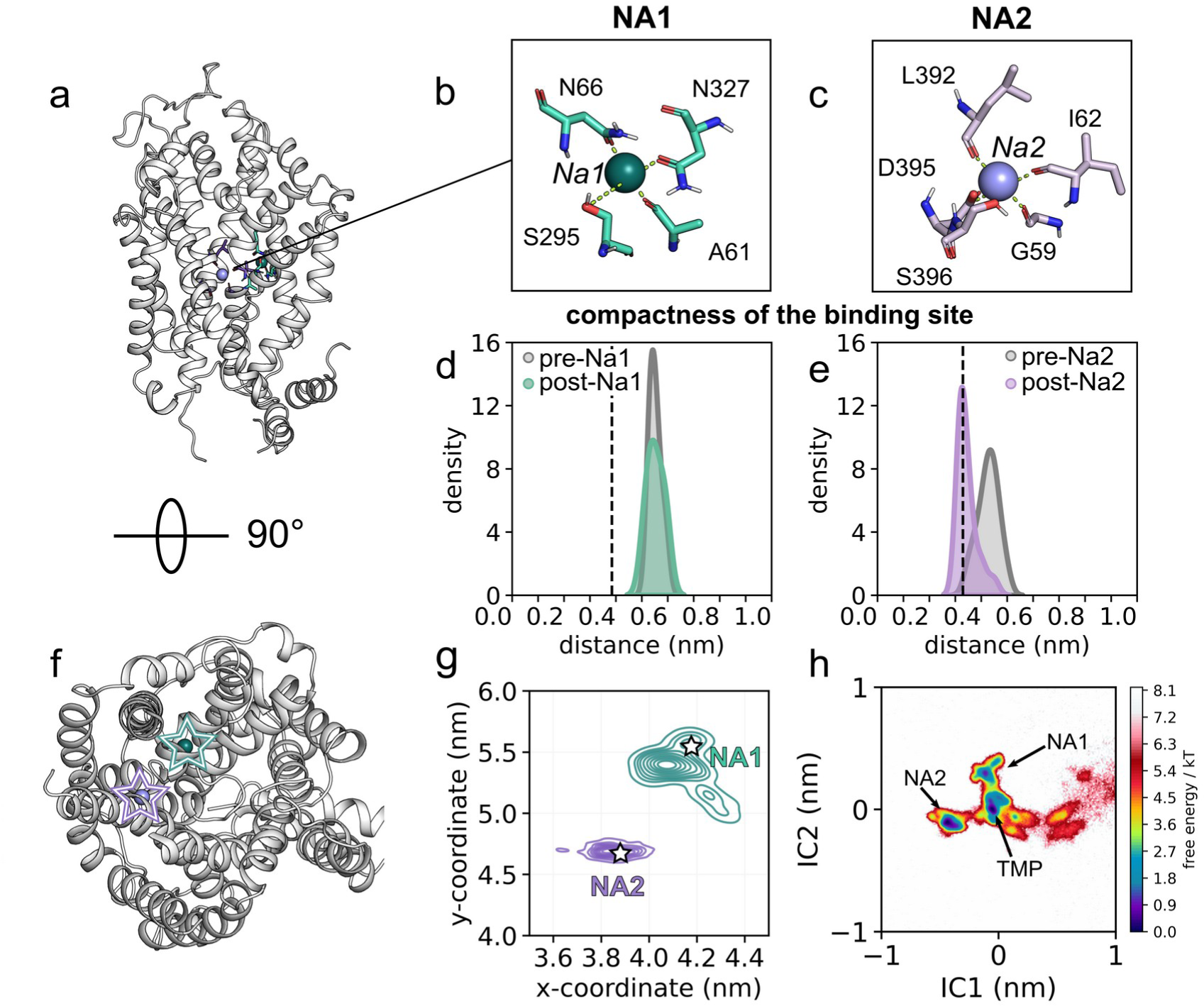
Short-range effect of sodium binding. **a)** GAT1 structure represented as a cartoon, highlighting the ion binding sites **b)** Visualisation of the sodium binding sites NA1 (green) and **c)** NA2 (purple). The dashed lines indicate the distances used for determining the compactness NA1 and NA2. **d)** Compactness of NA1 and **e)** NA2 before (in grey) and after sodium binding (coloured according to panels a and b) The vertical dashed lines indicate the measure in the cryo-EM structure (PDB ID: 7Y7W), where GAT is pictured in a holo inward-occluded conformation. **f)** Reference orientation of GAT1 as used for panel g. **g)** Top view projection of sodium ion positional density after binding to NA1 and NA2. The canonical binding positions of Na1 and Na2 are indicated by stars. **h)** Free energy landscape of sodium binding, where coordinates are projected along IC1 and IC2, respectively.

The dynamics of Na1 and Na2 differed (Figure 5g). While Na2 was firmly bound to NA2, Na1 remained dynamic. Some trajectories showed an unbinding as Na1 returned to the temporary side and in a few trajectories the sodium ion subsequently bound to NA2. The low binding stability of Na1 is consistent with the lack of an induced fit effect and indicates that interactions of Na1 with the NA1 are weak. We use time-lagged Independent Component Analysis (tICA) (Molgedey and Schuster, 1994; Naritomi and Fuchigami, 2013), which is a dimension reduction method based on the slowest motions, in combination with Markov State Modelling (Zwanzig, 1983) that allows for quantifying the transition probabilities between microstates to derive the underlying free surface as implemented in PyEMMA (Scherer et al., 2015). By applying this method, the simulation frames are first assigned to microstate to discretise the functionally important conformational space as defined by the dimension reduction method tICA. In a second step, a Markov State Model is derived from the transition probabilities between the microstates as observed in the simulations. This analysis allows to derive the free energy landscape of sodium translational motions and to project them into the subspace of the slowest two motions as identified by tICA. The results showed a deep energy minimum for NA2 and a shallow minimum for NA1 (Figure 5h). Consistent with the repeatedly observed unbinding of Na1 and the transient binding of sodium to the temporary site before reaching NA1 or NA2, this temporary site in the S1 shows a broad and shallow energy minimum. The energy barrier separating NA1 from the temporary site is small, while a larger energy barrier separates the NA2 from the temporary site, supporting the interpretation of weak binding of Na1 and strong binding of Na2. The free energy profile in the S1 is well described, with many transitions and a dense distribution of microstates. In contrast, the fast transition from the recruitment side to the S1 resulted in a sparse density of microstates, only a few transitions and thus in poor convergence. The relative free energy levels between the recruitment site and the S1 are therefore less certain.

### Na2 binding to the outward-facing conformation reduces internal transporter dynamics, thereby stabilising the closed inner gate and increasing structural integrity of TM5a

Next, we posed the question, if sodium binding to NA1 or NA2 would exert an allosteric effect on the overall transporter conformation and dynamics. We used principal component analysis (PCA) to identify the primary motions that occurred prior to and after sodium binding. PCA is a dimension reduction method that allows for identifying and extracting the motions with the largest amplitude in multidimensional data sets. We used PCA to project the dynamics of GAT1 onto the first two principal components (PCs) which represent the two motions with the largest amplitudes. Consistent with the lack of a compacting effect, we find that sodium binding to the NA1 did not have a global structural or dynamic effect on GAT1 (Figure S3). In contrast, binding of sodium to NA2 stabilised GAT1 in the outward-open conformation, which locally glued together TM1a and TM8. The projection of the two largest motions (PC1 and PC2) of GAT1 before and after sodium binding to NA2 (Figure 6a,b) showed that GAT1 samples multiple conformations in the absence of Na2, consistent with a dynamic transporter. Upon sodium binding, the trajectories showed convergence towards the outward-open conformation, confirming that Na2 stabilises GAT1 in the outward-open state (Szöllősi and Stockner, 2022). Projection of motions along PC1 as vectors on the TM helices of GAT1 (Figure 6c,d) showed that PC1 is mainly associated with motions of the bundle domain against the scaffold domain, whereby TM6a showing the largest amplitudes. In the absence of Na2, three basins can be observed along PC1, while after Na2 binding only one remains. In addition, PC1 captures motions of lower TM1a (residue 49 to 60) against TM5a (residue 242 to 245) and reduced mobility of the latter. Also PC2 describes motions of the bundle domain, though orthogonal to PC1. Helices with the largest amplitude are TM2, TM6 and TM12.

**Figure 6.**
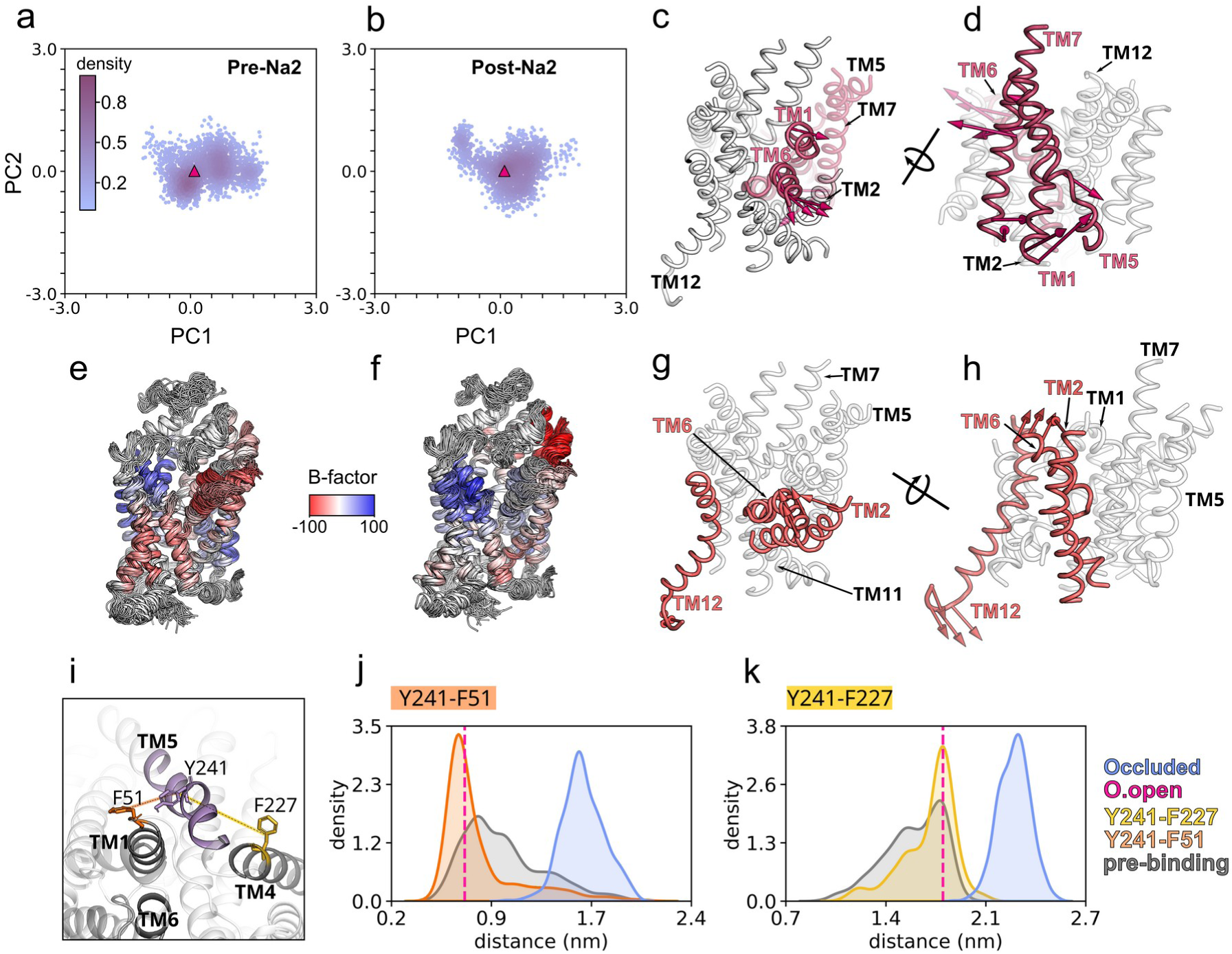
Global structural effects of sodium binding to NA2. Scatter plot of principal component 1 (PC1) against PC2 of all simulations that showed NA2 binding. Trajectories are divided into **a)** before and **b)** after Na2 binding. The pink triangle corresponds to the outward-open AlphaFold model of GAT1. All trajectories are projected to the same PC1 and PC2 axis. (**c,d**) Direction of motions along PC1 mapped as vectors on the TM helices of GAT1 **g,h)** Motions of GAT1 along PC2. (**e,f**) Using the same orientation as in panel c, superpositioning of 1 ns separated frames with the maximum likelihood method. Structures are coloured according to the degree of correlated motions along PC1 for frames **e)** before Na2 binding and **f)** after Na2 binding. **g)** Distribution of the distances Y241-F51 and Y241-F227 visualised in panel h, where values before Na2 binding are shown in grey, and after Na2 binding are coloured according to the legend. The vertical dashed lines indicate the distance measured in the cryo-EM structures of the GAT1 holo occluded and apo inward-open conformations (PDB ID: 7Y7W) (Zhu et al., 2023) and in the AlphaFold outward-open model **h**) The distances shown in panels g are indicated in orange between TM1a (F51) and TM5a (Y241), and in yellow between TM4 (F227) and TM5b (Y241)

This data indicates that stabilisation of GAT1 in a specific conformation by Na2 is moshttps://www.zotero.org/google-docs/?3bciiot likely a consequence of a reduction in bundle domain motions (Figure 6e,f). Similar to PCA, maximum likelihood structural alignment and correlation analysis showed that the largest motion of GAT1 prior to Na2 binding were concerted motions of the bundle domain relative to the scaffold domain. The motions of bundle and scaffold domains at the outer and the inner vestibule were anticorrelated with respect to each other, which is consistent with the rocking bundle model of SLC6 transporter motions. Consistent with the PCA analysis, at the cytosolic side the first part of TM5a showed large motions. Upon binding of Na2, the transporter dynamics were very different. Motions of the bundle domain relative to the scaffold domain were strongly diminished and breathing motions between the extracellular ends of TM5 and TM11 dominated. Na2 binding has therefore a long-range effect on the entire bundle domain, but also on TM5a, which contributes to the formation of the inner hydrophobic gate. Similar observations of a destabilisation of TM5a were linked to the inward-open conformation in both SERT (Coleman et al., 2019; Merkle et al., 2018) and LeuT, and were suggested to facilitate the creation of a permeation path from the intracellular side to the NA2 site. We have shown for SERT that destabilisation of TM5a is initiated by substrate occlusion, during which motions of TM5b follow the occluding movement of the bundle (Gradisch et al., 2024), while simultaneously, TM5a elongates as it remains attached to the scaffold domain. As a consequence, the helical secondary structure of TM5a destabilises and contributes to the opening of the inner gates by weakening of interaction to the bundle domain.

Na2 makes an important contribution to the stabilisation of the inner gate. To quantify the stability of the inner gate, we measured two orthogonal distances at the hydrophobic inner gate (Figure 6i): the contact across the hydrophobic inner gate between F51 (TM1a) and Y241 (TM6), as well as a measure of the length of TM5a by determining the distances between Y227 (TM4) as fixed reference point and Y248 on TM5a. Figure 6i,j shows distance distribution plots for the two measures. We found that the inner hydrophobic gate is dynamic in the absence of Na2, as GAT1 oscillates by 1 nm along the axis of inner gate opening (F51-Y241), indicating structural instability. The gate stabilised in the closed conformation in the presence of Na2, leading to a single main distance at ∼0.8 nm. Similar to the stability of the inner hydrophobic gate, TM5a (Figure 6g) showed a broader distribution in the absence of Na2. It condensed to a single main distance of ∼1.9 nm after Na2 binding, indicating that Na2 has a long-range effect on TM5a that leads to its stabilisation. Consistent with GAT1, movements of TM5a have been associated with opening of the inner gate at other members of the SLC6 transporter family (Coleman et al., 2019; Merkle et al., 2018). To confirm that transporter occlusion alone is sufficient to weaken the inner gate and to prime GAT1 for reaching the inward-open conformation (as observed for SERT (Gradisch *et al*., 2024)), we carried out five simulations (200 ns each) of the GABA-bound occluded conformation of GAT1 using the recently published occluded structure of *GAT1 (PDB ID: 7Y7W)* (Zhu *et al*., 2023). Applying the same measures (Figure 6j, k), the simulations showed that in the occluded conformation TM5a is extended by ∼0.5 nm with respect to the NA2-bound outward-open state and that TM1a separated from TM5a by ∼1.0 nm. The observed distributions for the GABA-bound occluded conformation of GAT1 are narrow, suggesting that complete opening of the inner gate is slow and might require an initial movement of Na2 to further destabilise the occluded state to promote a transition to the inward-open state.

### Role of NA1 in substrate binding

The behaviour of Na1 differs between the paralogue transporters SERT and GAT1. In SERT, Na1 binds tightly and the NA1 changes conformation according to an induced fit effect (Szöllősi and Stockner, 2021). A key difference between SERT and GAT1 is the Na1 coordinating carboxylate moiety (Figure 7b). In SERT, the carboxylate moiety is donated by the conserved aspartate residue D98, while in GAT1, the corresponding residue is a conserved glycine (G63) and, similar to LeuT (Yamashita et al., 2005), the carboxylate moiety of the substrate completes the Na1 (Figure 6c) coordination shell in GAT1.

**Figure 7.**
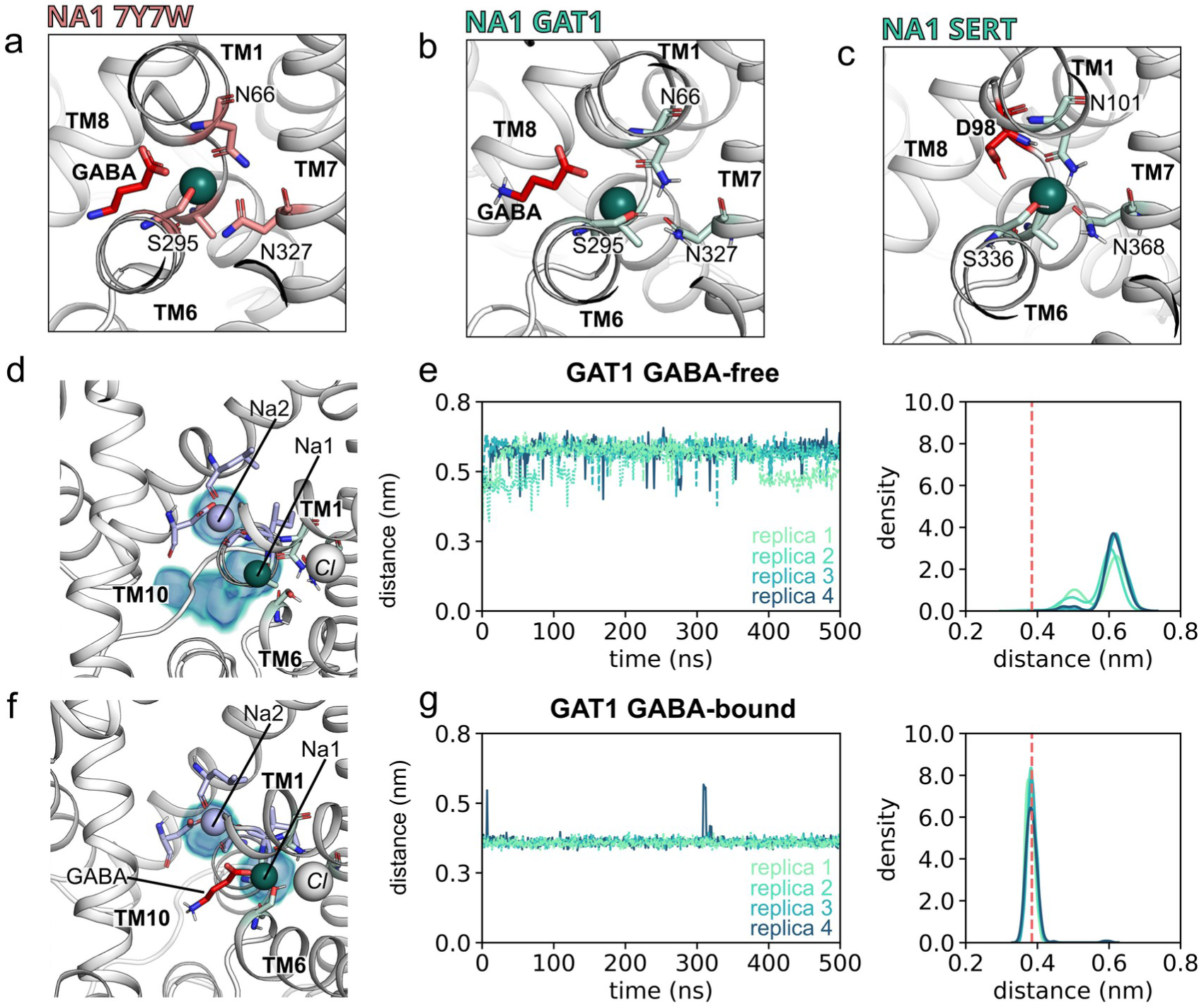
The substrate GABA completes the sodium binding site NA1. **a-c)** Comparison of the sodium binding site NA1 in **a)** the cryo-EM holo inward-occluded GAT1 (PDB ID: 7y7w (Zhu et al., 2023), **b)** AlphaFold holo outward-open GAT1, **c)** and SERT. The group donating the sodium-coordinating carboxylate moiety is GABA in GAT1 (a, b) and D98 in SERT c). **d)** Zoom onto S1 showing as isovolume densities the regions in which sodium ions Na1 and Na2 are found in the four 0.5 μs long simulations in the absence of GABA. **e**) Time course and averaged histogram of the compactness of NA1. The vertical dashed line highlights the measured NA1 compactness in the holo inward-occluded GAT1 cryo-EM structure (PDB ID: 7Y7W (Zhu et al., 2023) **f,g)** Same data as shown in panels d, and e, but in the presence of GABA.

To test this hypothesis, we docked GABA into sodium and chloride bound GAT1, carried out four independent 0.5 μs long simulations and compared the trajectories to four 0.5 μs long simulations in the absence of GABA (Figure 7d-g). Consistent with the sodium binding simulations, Na1 was unstable and temporarily unbound from the NA1 in the absence of GABA, moving to the temporary site, but remaining within the S1. The compactness of the NA1 (Figure 7e) was consistent with the compactness of the NA1 as observed in the sodium binding simulations (Figure 6c). Simulations carried out in the presence of GABA were strikingly different: Na1 remained stably bound and the NA1 was more compact. The compacting of the NA1 was reminiscent of an induced fit effect (Figure 7g). This data shows that the carboxylate moiety is essential for completing NA1 and is required to stabilise the Na1.

## Discussion

The importance of sodium for substrate transport has been characterised at the macroscopic and the kinetic level (Hilgemann and Lu, 1999; Iversen and Snyder, 1968; Mager et al., 1993) and showed that GABA uptake is energised by the downhill concentration gradient of the co-transported sodium (Kanner, 1978). Structural data of transporters from the SLC6 family have revealed the binding sites of substrate and co-transport ions and the main conformations of the transport cycle of GAT1 (Nayak et al., 2023; Zhu et al., 2023).

We confirmed through direct observations that NA2 functions to stabilise GAT1 in the outward-open conformation and showed that Na2 achieves this by immobilising the scaffold and bundle domains in relation to each other. Meanwhile, NA1 plays a crucial role in securing the GAT1 substrate by directly interacting with its carboxylate component, forming the essential connections that constitute the primary coordination environment of Na1.

Despite all this information, it is not completely clear how the sodium ions would be able to control the GABA transporter at the molecular level. The free energy landscape showed that NA2 binds sodium much stronger than NA1, and that the energy barrier connecting NA1 with a temporary site in the S1 is small. This is reflected in the dynamics of sodium ions. Simulations showed a clear order for sodium binding: the first stably bound sodium enters the NA2 and stabilises the outward-open conformation, thereby priming GAT1 for substrate binding. Once bound to the NA2, we did not observe any indication for unbinding of Na2 during the time course of the simulations. Sodium ions, which initially interacted with NA1, remained mobile, eventually returning to the temporary site in the S1 and possibly bound to the NA2 (if empty). The second sodium ion that is electrostatically attracted to the S1 remains in a loose binding state as it keeps oscillating between the NA1 and the temporary site, because the NA1 remains incomplete until a substrate reaches the S1. Access to the S1 remains open in the outward-open state, irrespective of the presence of 0, 1, or 2 sodium ions. Importantly, in the outward-open state, the sidechain of F294 does not rotate into the vestibule to obstruct access to the S1. In analogy to SERT (Gradisch *et al*., 2022) we can assume that the side chain of F294 reorients towards the vestibule to form a closed outer hydrophobic lid only once approaching the occluded state. The role of Na2 is to stabilise GAT1 in the outward-open conformation. It was shown for LeuT that the transporter isomerizes between the inward-facing and the outward-facing state with comparable populations for both conformations in the absence of sodium ions, while the presence of sodium stabilises the outward-facing conformation (Zhao et al., 2011). A similar conformational effect upon sodium binding was observed for human SLC6 transporters (Schicker *et al*., 2012; Tavoulari *et al*., 2016; Zhang *et al*., 2018; Szöllősi and Stockner, 2022), because the NA2 is located between the scaffold domain and the bundle domain, specifically between TM1a and TM8 (Krishnamurthy and Gouaux, 2012; Yamashita et al., 2005). NA2 becomes disrupted by a transition to the inward-facing state. These data remain inconclusive with respect to the functionally important question, if Na2 would stabilise the NA2 and thus the transporter in the outward-open state or if the outward-open state would be formed by other interactions and thereby creating the sodium binding competent NA2. Our data showed that it is Na2 binding that stabilises GAT1. We observed that GAT1 is dynamic in the outward-open state and showed large bundle domain motions on both the extracellular and the intracellular side in the absence of Na2. This is consistent with a dynamic state reminiscent of the apo state of LeuT without conformational preference (Zhao et al., 2011). We also observed motions at the inner gate, specifically between TM5a and the bundle domain helices TM1a and TM6b, which indicate that Na2 binding plays a long-range stabilising role for the inner gate and for TM5a. A closed hydrophobic inner gate is necessary for stabilising the outward-facing state, while it has to open for reaching the inward-open state. We found a dynamic inner gate in the absence of Na2, as we measured broad distributions of distances across the inner gate. A long range effect of Na2 binding resulted in the disappearance of these dynamic motions, leading to the stabilisation of a sealed inner gate. We find that a long range allosteric effect of Na2 binding is a stabilisation of the conformation of TM5a, reminiscent of the observations in the homologous transporters LeuT and SERT (Merkle *et al*., 2018; Coleman *et al*., 2019; Gradisch *et al*., 2024), which showed an unwinding and an extension of TM5a in the transition from the outward-facing to the inward-facing state. By connecting the bundle domain to the scaffold domain, TM5 follows the motions associated with the transitions of the transport cycle, whereby TM5a is attached to the scaffold domain and TM5b is linked to the bundle domain. The motions of the bundle domain, responsible for this property, were quenched after sodium binding to NA2, showing how Na2 stabilises the outward-open state, and distances are extended once reaching the occluded state. The conformation stabilising effect of Na2 is achieved by an induced fit effect. Analysis of the local environment of the sodium coordinating oxygen atoms at the NA2 showed a clear structural change upon sodium binding that brought the NA2 exposed residues of TM1a and TM8 into a defined distance that is smaller than these residues would be in the absence of Na2. This local structural change fixes these two helices relative to each other and thereby prevents bundle domain motions relative to the scaffold domain.

The NA1 site has the role of stabilising the bound substrate in the S1. The apparent affinity of GABA for GAT1 increases strongly in the presence of sodium (Radian and Kanner, 1983), indicating cooperative binding. Here we show that binding of Na1 and of GABA is synergistic, as GABA stabilises Na1 in the NA1 and Na1 stabilises GABA in the S1. We observed by direct simulations that sodium oscillated between the NA1 and a temporary site in the S1 without induced fit conformational changes of NA1 in the absence of substrate. In contrast, in the presence of GABA, sodium is firmly bound to the NA1, the carboxylate moiety of GABA completes the first coordination shell of Na1 and NA1 compacts. The same position for the carboxylate was found in the occluded structure of GAT1 (Zhu et al., 2023). As yet no structural information on Na-bound GAT1 in the absence of GABA is available, but different coordination of Na1 in LeuT were observed in the absence and presence of the coordinating carboxyl group of the amino acid substrate (Krishnamurthy and Gouaux, 2012), it is likely that this difference in Na1 coordination is similar in GAT1. The carboxyl group of GABA overlaps with the carboxylate of D98 in the paralogues serotonin transporter SERT, where sodium binds strongly in the absence of substrate and the NA1 compacts due to an induced fit effect (Szöllősi and Stockner, 2021). The difference between GAT1 and SERT is therefore that strong binding of Na1 and the NA1 condensation in GAT1 is GABA dependent, while in the SERT it is substrate independent. From this data we can infer that the strong electrostatic interactions between the negatively charged carboxylate of GABA and the positive charge of sodium as well as the condensation of NA1 due to the induced fit effect contribute cooperatively to the affinity of GABA to GAT1. GABA adopts a binding mode in which the carboxyl group completes the coordination of sodium in NA1 but is also stabilised by a hydrogen bond to Y140 of the scaffold domain. It is tempting to speculate that this interaction relay from Na1 (bundle domain associated), GABA to Y140 (scaffold domain) could be the switch that triggers GAT1 occlusion and thus substrate uptake reminiscent of the mechanism described by Gradisch et al. (Gradisch et al., 2022).

For the simulations we used the AlphaFold model of outward-facing GAT1, as no outward-facing structure of GAT1 is available. The initial use of SERT based outward-open homology models resulted in GAT1 structures in which the inner hydrophobic gate was not sealed, allowing ions to penetrate. Thus, these models were not further used. A sequence comparison showed that this property is a consequence of a set of smaller side chains in GAT1 compared to SERT at the interface between the scaffold and the bundle domain. The outer vestibule of the AlphaFold model is somewhat less open as compared to the SERT based model, but PC1 and PC2 (Figure 6a,b) show that the AlphaFold starting structure is inside the highly populated basin of conformations accessed by GAT1 during simulations, confirming that the AlphaFold model represents a valid starting structure for the outward-open GAT1.

Sodium binding and substrate transport by GAT1 are accelerated by a sodium recruitment site. Our simulations showed consistently a region of high sodium density at the entry to the outer vestibule. The sodium recruitment site consists of two negatively charged residues located at the beginning of TM6a that electrostatically attracts sodium ions. *In-silico* mutations of these residues resulted in the disappearance of enhanced sodium density at the sodium recruitment site and slowed binding kinetics (slower on rate) for sodium entering the S1 (Figure 1). *In-vitro* experiments confirmed these predictions (Figure 3-4) by showing a reduced sodium dependency of GABA uptake and a lower Vmax, while the Km of GABA uptake was almost unchanged. These data indicate that the role of the sodium recruiting site is to increase the probability of sodium ions to enter the outer vestibule for reaching the sodium binding sites NA1 and NA2. The negative electrostatic potential at the sodium recruitment site attracts positively charged sodium ions and locally increased sodium concentration at the entry into the outer vestibule. In contrast, our data showed that EL4, and in particular EL4a and its only negatively charged residue D355, do not play a prominent role. This is most likely a consequence of the high number of positively charged residues (R69,Y76, K350, R351) surrounded D355, which reduce the local density of sodium ions at the EL4a by electrostatic repulsion. We did not observe tight binding to the recruitment site, but instead observed that sodium ions eventually unbind. This is kinetically important, because it increases the on rate, as the sodium ion can leave either towards the extracellular solution or it is efficiently steered towards the S1 by electrostatic attraction.

## Material and Methods

### Molecular Dynamics simulations

In the absence of a crystal or cryo-EM structure of GAT1 in the outward-facing conformation, we selected the AlphaFold model to investigate sodium binding (Jumper et al., 2021; Varadi et al., 2022). The bound-chloride ion was added to the GAT1 model using as a reference the outward-open human SERT crystal structure (PDB ID: 5I71; (Coleman et al., 2016). The generated system was converted from full-atom into a coarse-grained representation using the MARTINI force field (de Jong et al., 2013; Monticelli et al., 2008; Wassenaar et al., 2015), with a membrane composition of 1-palmitoyl-2-oleoyl phosphatidylcholine (POPC): cholesterol containing membrane (POPC:CHOL 70:30 mol%; (van Meer, 1998)), and solvated in water with 150 mM NaCl. The membrane was equilibrated after a 1 μs long coarse-grained simulation, while restraining the protein. The coarse-grained system was then converted to an all-atom representation (Ingólfsson et al., 2014) in which the transporter was replaced by the original GAT1 model to avoid spurious structural imprecisions induced by the double coordinate conversion protocol. Possible atom overlaps between the reinserted protein and the equilibrated membrane were relaxed using the membed procedure (Wolf et al., 2010) as previously described (Szöllősi and Stockner, 2022). We used the amber ff99SB-ILDN force field (Lindorff-Larsen et al., 2010) to describe GAT1, ions and the solvent, and Slipid (Jämbeck et al., 2013; Jämbeck and Lyubartsev, 2013) for POPC and cholesterol. Previously reported structural analysis of the orthologous SERT structure (Coleman et al., 2016) suggested that the residue E467 in GAT1 is protonated. All simulations were carried out with GROMACS version 2019.2 (Abraham et al., 2015). Three replicas of the final assembled system were energy-minimised and equilibrated in four steps of 2.5 ns, each by stepwise releasing the position restraints (1,000, 100, 10, 1 kJ/mol/nm) that are active on the Cα atoms and the bound chloride ion (Goda et al., 2020; Sohail et al., 2016). Per each trajectory, the production run was carried for 500 ns. The temperature was maintained at 310 K using the v-rescale (τ = 0.5 ps) thermostat (Bussi et al., 2007), while separately coupling protein, membrane, and solvent. The pressure was maintained at 1 bar using the Parrinello-Rahman barostat (Parrinello and Rahman, 1981) in a semi-isotropic manner and applying a coupling constant of 20.1 ps. Long-range electrostatic interactions were described using the smooth particle mesh Ewald method (Darden et al., 1993) applying a cutoff of 0.9 nm. The van der Waals interactions were described using the Lennard Jones potentials applying a cutoff of 0.9 nm. Long-range corrections for energy and pressure were applied. Coordinates of all atoms were recorded every 5 ps. The complete set of parameters of the production run can be found in the Supplementary Materials. The simulations of the holo inward-occluded GAT1 (PDB: 7Y7W) were conducted by following the same protocol, and comprised 4 repeats, each of which being 200 ns long.

The trajectories that were initiated from the AlphaFold model served as parent trajectories and were used to extract starting structures for the sodium binding simulations. From each replica, 20 equally spaced frames were extracted every 10 ns of the 200 ns long trajectories, which then served as starting structures after removal of sodium ions that might be present in the substrate binding pocket S1. Each of these 60 systems was simulated for 200 ns with the same simulation parameters as the parent trajectories. The same starting systems were used for the double mutant (D281A-E283A) systems and the same simulation parameters were applied. Mutants were created using the Pymol Mutagenesis Wizard tool (Schrödinger, LLC, Warren DeLano, 2020).

To generate the trajectory for investigating the role of sodium Na1, we extracted a frame from the parent trajectory in which Na2 was already bound. For the GABA-free systems, we placed the second sodium into the temporary site of the S1 as observed in the sodium binding simulations; for the GABA-bound systems we placed GABA in the binding site of GAT1, in a similar way as leucine was observed in the LeuT crystal structure (Yamashita et al., 2005). Figures and statistical analyses were generated by the GROMACS package, R, and python scripts using the MD Analysis package, v0.19.2 (Gowers et al., 2016; Michaud-Agrawal et al., 2011). For visualisation VMD (Humphrey et al., 1996) v1.9.3 and Pymol v1.8.4 (Schrödinger, LLC, Warren DeLano, 2020) were used.

### Markov State Modelling

The Markov State Models (MSMs) were built with the pyEMMA python package version 2.5.9 (Scherer et al., 2015). To begin, we defined the reaction coordinates employing time-lagged Independent Component Analysis (tICA) (Molgedey and Schuster, 1994; Naritomi and Fuchigami, 2013), an advanced linear transformation technique design to maximise the autocorrelation of molecular descriptors. This facilitates the identification of slowest kinetic modes within the system, i.e., the reaction coordinates. The input dataset for tICA comprised the precise coordinates (x, y and z) of the sodium ions undergoing binding events. The dimensionality reduction was performed using a time lag of 1 ns which allowed for a complete exploration of the temporal dynamics. Subsequently, the trajectories were discretized into 500 cluster centres by applying the k-mean clustering algorithm. Finally, the MSMs were estimated using a time lag of 25 ns, the time point at which we observed convergence of the implied timescales. The Markovian behaviour was assured using the Chapman-Kolmogorov (CK) test (Figure S4). The free energy values were estimated using the MSMs based on the relationship derived from the stationary distribution (π) values, as obtained from the transition probability matrix. This relationship is represented mathematically as GSj=-K_b_Tln (sum(πj)). Where j denotes the MSM stationary weight of the j^th^ microstate. G, S, K_b_ and T represent the Gibbs free energy, entropy, Boltzmann constant and temperature, respectively. To visualise and interpret the free energy landscape associated with sodium binding, we plotted the free energy values into the tICA subspace, specifically along the first and second slowest motions, IC1 and IC2.

### Cell lines and cell culture

An N-terminal eYFP tagged human GAT1 wild-type plasmid was used. The Agilent QuikChange Primer Design Program and the QuikChange II Site-Directed Mutagenesis Kit (Agilent Technologies, Santa Clara, USA) were utilized to incorporate point mutations into the coding sequence of human GAT1. After construct verification (LGC Genomics, Berlin, Germany) double sorted polyclonal HEK293 cell lines, stably and consistently expressing hGAT1 wild-type, the single mutants D281A and E283A as well as the double alanine variant (D281A-E283A) were established, as previously described (Gradisch *et al*., 2024). Therefore, raw HEK cells were transfected with the respective plasmid by applying the jetPRIME transfection protocol (VWR International GmBH, Vienna, Austria). Adding 100 µl geneticin (G418, 50 mg x ml^−1^) ensured high selection pressure which was maintained for at least 14 days before cells were sorted by using fluorescence-activated cell sorting (FACS). Subsequently cells were grown for 1 week followed by a second round of FACS sorting. Cells stably expressing GAT1 were grown in 10 cm cell culture dishes (Greiner) with Dubleco’s Modified Eagle Media (DMEM, Sigma-Aldrich, St. Louis, USA) supplemented with 10% heat-inactivated Fetal Bovine Serum (FBS, Sigma Aldrich), penicillin (1 U x ml^−1^, Sigma Aldrich), streptomycin (100 µg x ml^−1^, Sigma Aldrich) and geneticin (50 µg x ml^−1^, Sigma Aldrich) at 37 °C and 5 % CO2 in an incubator. Regular tests for mycoplasma were performed by 4′,6-diamidino-2-phenylindole staining.

### Confocal microscopy

A day prior to live confocal imaging the cells were seeded onto poly-D-lysine (PDL)-coated 35 mm glass-bottom dishes (Cellvis, Sunnyvale, California, USA) at a density of 0.2 x 10^6^ cells per 2 ml). After aspiration of the culture medium, cell membranes were stained with a 0.4% Trypan blue containing solution (Sigma) for 10 minutes. Subsequently, cells were washed several times with Krebs-HEPES buffer (KHP, 10 mM HEPES, 120 mM NaCl, 3 mM KCl, 2 mM CaCl2, 2 mM MgCl2, 2 mM glucose monohydrate, pH 7.3) and kept on KHB throughout the imaging process. Cell imaging was performed using a Nikon A1R+ laser scanning confocal microscope equipped with a 60x NA1.4 oil immersion objective (Nikon, Vienna, Austria). To visualise Trypan blue and eYFP fluorescence, laser lines at 561 nm and 488 nm were employed, respectively. Prior light detection by a GaAsP PMT detector, emitted light was filtered with a 595/50 nm (trypan blue) and 525/50 (eYFP) nm emission filter. Image analysis was performed by using Fiji ImageJ 1.53c. Trypan blue staining confined both regions of interest (ROI) used to measure surface expression as well as cytosolic retained transporters. Both measurements were defined by mean fluorescence over the confined ROI.

### Radiotracer Uptake assays

Cells were seeded (∼0.05 x 10^6^ cells / 0.2 ml / well) onto PDL-coated 96-well plates a day before the experiment. After removal of the culture medium cells were kept on KHP throughout the experiment. Concentration-dependent uptake was determined by applying solutions containing 50 nM [^3^H]GABA and increasing concentrations of GABA (0.15 - 59.95 µM) for 3 minutes at room temperature. Thereafter, cells were washed with 500 µl ice-cold KHB to terminate the uptake and subsequently lysed by adding 200 µl Ultima Gold Scintillation Cocktail (Sigma).

In case of determining the sodium-dependence of GAT1-mediated [^3^H]GABA uptake, cells were kept in KHP with varying concentrations of sodium (0 mM - 200 mM). To compensate for changes in osmolarity N-methyl D-glucamine (NMDG)-chloride was added accordingly (200 mM - 0 mM). Uptake solutions (KHB with varying concentrations of sodium and NDMG-Cl) containing 50 nM [^3^H]GABA were added to the cells for 3 minutes, followed by aspiration and a washing step with 500 µl ice-cold KHB. Cell lysis was achieved by adding 200 µl of Ultima Gold Scintillation Cocktail. Non-specific uptake was determined in presence of 10 µM tiagabine. A ß-scintillation counter (Perkin Elmer, Waltham, USA) was used to quantify the uptake of [^3^H]GABA.

### Whole-cell patch clamp

24 hours prior the experiment cells were seeded at very low density onto PDL-coated 35 mm dishes. hGAT1-mediated and substrate induced currents were recorded in whole-cell configuration at a holding-potential of −60 mV. Throughout the experiments cells were maintained in regular external solution (bath solution) containing: 140 mM NaCl, 20 mM D-glucose, 10 mM HEPES, 3 mM KCl, 2.5 mM CaCl_2_, 2 mM MgCl_2_, pH adjusted to 7.4 with NaOH. The patch pipette harbours regular internal solution that comprises: 133 mM K^+^-gluconate, 10 mM HEPES, 10 mM EGTA, 5.9 mM NaCl, 1 m CaCl_2_, 0.7 mM MgCl_2_, pH adjusted to 7.2 with KOH. An 8 tube ALA perfusion manifold (NPI Electronic GmbH, Germany) and a DAD-12 superfusion system (Adams & List, Westbury, NY) ensured fast drug perfusion onto the cell and complete solution exchange surrounding the cell within ∼100 ms. GABA was applied for 5 seconds. After passive holding currents were subtracted, the traces were filtered by applying a 100 Hz digital Gaussian low-pass filter as well as a 50 Hz harmonics filter. Clampfit 10.2 was used to analyse the current amplitudes. Data analysis was performed by using GraphPad Prism version 9.4.1. 11 independent cells per condition were recorded.

### Data and statistical analysis

Experimentally acquired data were analysed and plotted with GraphPad Prism 10.1.0 (GraphPad Software Inc., San Diego, USA). Michaelis Menten kinetics (Km, Vmax) were determined solving the following equation: *Y* = Vmax*X/(Km + X). EC_50_ values deduced from electrophysiological recordings were determined as follows: Y=Bottom + (Top-Bottom)/(1+10^((LogEC50-X))). The affinity for sodium (EC_50_) was calculated by solving Y=Bottom + (X^Hillslope)*(Top-Bottom)/(X^HillSlope + EC50^HillSlope). All data are from at least five biologically independent experiments (*n* ≥ 5), measured in triplicates and reported as mean ± SD. In case of electrophysiological measurements 6 individual cells were recorded and plotted as mean ± SD. Statistics were conducted by applying the one-way ANOVA test followed by a Dunnett’s post hoc multiple comparison vs. wild-type. Statistical significance was denoted as follows: *=P < 0.05, **=P < 0.01, ***P < 0.001, ****P < 0.0001.

## Data availability

The data that support the findings of this study are available on the ZENODO open data repository (DOI:10.5281/zenodo.10686813). The dataset includes all starting and end structures of all simulations and all dataset used for creating the figures.

## Author Contributions

Erika Lazzarin: Conceptualization, Data curation, Formal analysis, Investigation, Methodology, Writing, Original Draft Preparation; Ralph Gradisch: Conceptualization, Data curation, Formal analysis, Investigation, Review & Editing; Sophie M.C. Skopec: Investigation, Formal analysis, Methodology; Leticia Alves da Silva: Investigation, Methodology, Software; Marko Roblek: Investigation, Formal analysis, Methodology; Chiara Sebastianelli-Schoditsch: Investigation, Formal analysis, Methodology; Dániel Szöllősi: Methodology, Software, Writing, Review & Editing; Julian Maier, Investigation, Writing, Review & Editing; Sonja Sucic: Funding acquisition, Resources, Writing, Review & Editing; Baruch I. Kanner: Writing, Review & Editing; Harald H Sitte: Resources, Supervision, Funding acquisition, Writing, Review & Editing; Thomas Stockner: Writing, Conceptualization, Project Administration, Resources, Supervision, Original Draft Preparation, Review & Editing

## References

Abraham, M.J., Murtola, T., Schulz, R., Páll, S., Smith, J.C., Hess, B., Lindahl, E., 2015. GROMACS: High performance molecular simulations through multi-level parallelism from laptops to supercomputers. SoftwareX 1–2, 19–25. 10.1016/j.softx.2015.06.001

Bhat, S., El-Kasaby, A., Freissmuth, M., Sucic, S., 2021. Functional and Biochemical Consequences of Disease Variants in Neurotransmitter Transporters: A Special Emphasis on Folding and Trafficking Deficits. Pharmacol. Ther. 222, 107785. 10.1016/j.pharmthera.2020.107785

Bhatt, M., Gauthier-Manuel, L., Lazzarin, E., Zerlotti, R., Ziegler, C., Bazzone, A., Stockner, T., Bossi, E., 2023. A comparative review on the well-studied GAT1 and the understudied BGT-1 in the brain. Front. Physiol. 14.

Bicho, A., Grewer, C., 2005. Rapid substrate-induced charge movements of the GABA transporter GAT1. Biophys. J. 89, 211–231. 10.1529/biophysj.105.061002

Bussi, G., Donadio, D., Parrinello, M., 2007. Canonical sampling through velocity rescaling. J. Chem. Phys. 126, 014101. 10.1063/1.2408420

Chen, N.-H., Reith, M.E.A., Quick, M.W., 2004. Synaptic uptake and beyond: the sodium- and chloride-dependent neurotransmitter transporter family SLC6. Pflugers Arch. 447, 519–531. 10.1007/s00424-003-1064-5

Coleman, J.A., Green, E.M., Gouaux, E., 2016. X-ray structures and mechanism of the human serotonin transporter. Nature 532, 334–339. 10.1038/nature17629

Coleman, J.A., Yang, D., Zhao, Z., Wen, P.-C., Yoshioka, C., Tajkhorshid, E., Gouaux, E., 2019. Serotonin transporter–ibogaine complexes illuminate mechanisms of inhibition and transport. Nature 569, 141–145. 10.1038/s41586-019-1135-1

Darden, T., York, D., Pedersen, L., 1993. Particle mesh Ewald: An N⋅log(N) method for Ewald sums in large systems. J. Chem. Phys. 98, 10089–10092. 10.1063/1.464397

de Jong, D.H., Singh, G., Bennett, W.F.D., Arnarez, C., Wassenaar, T.A., Schäfer, L.V., Periole, X., Tieleman, D.P., Marrink, S.J., 2013. Improved Parameters for the Martini Coarse-Grained Protein Force Field. J. Chem. Theory Comput. 9, 687–697. 10.1021/ct300646g

Fischer, F.P., Kasture, A.S., Hummel, T., Sucic, S., 2022. Molecular and Clinical Repercussions of GABA Transporter 1 Variants Gone Amiss: Links to Epilepsy and Developmental Spectrum Disorders. Front. Mol. Biosci. 9.

Forrest, L.R., Zhang, Y.-W., Jacobs, M.T., Gesmonde, J., Xie, L., Honig, B.H., Rudnick, G., 2008. Mechanism for alternating access in neurotransmitter transporters. Proc. Natl. Acad. Sci. 105, 10338–10343. 10.1073/pnas.0804659105

Ghit, A., Assal, D., Al-Shami, A.S., Hussein, D.E.E., 2021. GABAA receptors: structure, function, pharmacology, and related disorders. J. Genet. Eng. Biotechnol. 19, 123. 10.1186/s43141-021-00224-0

Goda, K., Dönmez-Cakil, Y., Tarapcsák, S., Szalóki, G., Szöllősi, D., Parveen, Z., Türk, D., Szakács, G., Chiba, P., Stockner, T., 2020. Human ABCB1 with an ABCB11-like degenerate nucleotide binding site maintains transport activity by avoiding nucleotide occlusion. PLOS Genet. 16, e1009016. 10.1371/journal.pgen.1009016

Gowers, R., Linke, M., Barnoud, J., Reddy, T., Melo, M., Seyler, S., Domański, J., Dotson, D., Buchoux, S., Kenney, I., Beckstein, O., 2016. MDAnalysis: A Python Package for the Rapid Analysis of Molecular Dynamics Simulations. Presented at the Python in Science Conference, Austin, Texas, pp. 98–105. 10.25080/Majora-629e541a-00e

Gradisch, R., Schlögl, K., Lazzarin, E., Niello, M., Maier, J., Mayer, F.P., Alves da Silva, L., Skopec, S.M.C., Blakely, R.D., Sitte, H.H., Mihovilovic, M.D., Stockner, T., 2024. Ligand coupling mechanism of the human serotonin transporter differentiates substrates from inhibitors. Nat. Commun. 15, 417. 10.1038/s41467-023-44637-6

Gradisch, R., Szöllősi, D., Niello, M., Lazzarin, E., Sitte, H.H., Stockner, T., 2022. Occlusion of the human serotonin transporter is mediated by serotonin-induced conformational changes in the bundle domain. J. Biol. Chem. 298, 101613. 10.1016/j.jbc.2022.101613

Guastella, J., Nelson, N., Nelson, H., Czyzyk, L., Keynan, S., Miedel, M.C., Davidson, N., Lester, H.A., Kanner, B.I., 1990. Cloning and Expression of a Rat Brain GABA Transporter. Science 249, 1303–1306. 10.1126/science.1975955

Hasenhuetl, P.S., Freissmuth, M., Sandtner, W., 2016. Electrogenic Binding of Intracellular Cations Defines a Kinetic Decision Point in the Transport Cycle of the Human Serotonin Transporter. J. Biol. Chem. 291, 25864–25876. 10.1074/jbc.M116.753319

Hilgemann, D.W., Lu, C.-C., 1999. Gat1 (Gaba:Na+:Cl−) Cotransport Function: Database Reconstruction with an Alternating Access Model. J. Gen. Physiol. 114, 459–476. 10.1085/jgp.114.3.459

Humphrey, W., Dalke, A., Schulten, K., 1996. VMD: Visual molecular dynamics. J. Mol. Graph. 14, 33–38. 10.1016/0263-7855(96)00018-5

Ingólfsson, H.I., Melo, M.N., van Eerden, F.J., Arnarez, C., Lopez, C.A., Wassenaar, T.A., Periole, X., de Vries, A.H., Tieleman, D.P., Marrink, S.J., 2014. Lipid Organization of the Plasma Membrane. J. Am. Chem. Soc. 136, 14554–14559. 10.1021/ja507832e

Iversen, L.L., Snyder, S.H., 1968. Synaptosomes: Different Populations storing Catecholamines and Gamma-aminobutyric Acid in Homogenates of Rat Brain. Nature 220, 796–798. 10.1038/220796a0

Jämbeck, J.P.M., Lyubartsev, A.P., 2013. Another Piece of the Membrane Puzzle: Extending Slipids Further. J. Chem. Theory Comput. 9, 774–784. 10.1021/ct300777p

Jämbeck, J.P.M., Mocci, F., Lyubartsev, A.P., Laaksonen, A., 2013. Partial atomic charges and their impact on the free energy of solvation. J. Comput. Chem. 34, 187–197. 10.1002/jcc.23117

Jumper, J., Evans, R., Pritzel, A., Green, T., Figurnov, M., Ronneberger, O., Tunyasuvunakool, K., Bates, R., Žídek, A., Potapenko, A., Bridgland, A., Meyer, C., Kohl, S.A.A., Ballard, A.J., Cowie, A., Romera-Paredes, B., Nikolov, S., Jain, R., Adler, J., Back, T., Petersen, S., Reiman, D., Clancy, E., Zielinski, M., Steinegger, M., Pacholska, M., Berghammer, T., Bodenstein, S., Silver, D., Vinyals, O., Senior, A.W., Kavukcuoglu, K., Kohli, P., Hassabis, D., 2021. Highly accurate protein structure prediction with AlphaFold. Nature 596, 583–589. 10.1038/s41586-021-03819-2

Kanner, B.I., 1978. Active transport of γ-aminobutyric acid by membrane vesicles isolated from rat brain. Biochemistry 17, 1207–1211. 10.1021/bi00600a011

Kasture, A.S., Fischer, F.P., Kunert, L., Burger, M.L., Burgstaller, A.C., El-Kasaby, A., Hummel, T., Sucic, S., 2023. Drosophila melanogaster as a model for unraveling unique molecular features of epilepsy elicited by human GABA transporter 1 variants. Front. Neurosci. 16.

Keynan, S., Kanner, B.I., 1988. gamma-Aminobutyric acid transport in reconstituted preparations from rat brain: coupled sodium and chloride fluxes. Biochemistry 27, 12–17. 10.1021/bi00401a003

Kortagere, S., Fontana, A.C.K., Rose, D.R., Mortensen, O.V., 2013. Identification of an allosteric modulator of the serotonin transporter with novel mechanism of action. Neuropharmacology 72, 282–290. 10.1016/j.neuropharm.2013.04.026

Koshland JR., D.E., 1959. Enzyme flexibility and enzyme action. J. Cell. Comp. Physiol. 54, 245–258. 10.1002/jcp.1030540420

Krishnamurthy, H., Gouaux, E., 2012. X-ray structures of LeuT in substrate-free outward-open and apo inward-open states. Nature 481, 469–474. 10.1038/nature10737

Kristensen, A.S., Andersen, J., Jørgensen, T.N., Sørensen, L., Eriksen, J., Loland, C.J., Strømgaard, K., Gether, U., 2011. SLC6 neurotransmitter transporters: structure, function, and regulation. Pharmacol. Rev. 63, 585–640. 10.1124/pr.108.000869

Krnjević, K., Schwartz, S., 1967. The action of γ-Aminobutyric acid on cortical neurones. Exp. Brain Res. 3, 320–336. 10.1007/BF00237558

Lindorff-Larsen, K., Piana, S., Palmo, K., Maragakis, P., Klepeis, J.L., Dror, R.O., Shaw, D.E., 2010. Improved side-chain torsion potentials for the Amber ff99SB protein force field. Proteins 78, 1950–1958. 10.1002/prot.22711

Mager, S., Naeve, J., Quick, M., Labarca, C., Davidson, N., Lester, H.A., 1993. Steady states, charge movements, and rates for a cloned GABA transporter expressed in Xenopus oocytes. Neuron 10, 177–188. 10.1016/0896-6273(93)90309-F

Merkle, P.S., Gotfryd, K., Cuendet, M.A., Leth-Espensen, K.Z., Gether, U., Loland, C.J., Rand, K.D., 2018. Substrate-modulated unwinding of transmembrane helices in the NSS transporter LeuT. Sci. Adv. 4, eaar6179. 10.1126/sciadv.aar6179

Michaud-Agrawal, N., Denning, E.J., Woolf, T.B., Beckstein, O., 2011. MDAnalysis: A toolkit for the analysis of molecular dynamics simulations. J. Comput. Chem. 32, 2319–2327. 10.1002/jcc.21787

Molgedey, L., Schuster, H.G., 1994. Separation of a mixture of independent signals using time delayed correlations. Phys. Rev. Lett. 72, 3634–3637. 10.1103/PhysRevLett.72.3634

Monod, J., Wyman, J., Changeux, J.-P., 1965. On the nature of allosteric transitions: A plausible model. J. Mol. Biol. 12, 88–118. 10.1016/S0022-2836(65)80285-6

Monticelli, L., Kandasamy, S.K., Periole, X., Larson, R.G., Tieleman, D.P., Marrink, S.-J., 2008. The MARTINI Coarse-Grained Force Field: Extension to Proteins. J. Chem. Theory Comput. 4, 819–834. 10.1021/ct700324x

Motiwala, Z., Aduri, N.G., Shaye, H., Han, G.W., Lam, J.H., Katritch, V., Cherezov, V., Gati, C., 2022. Structural basis of GABA reuptake inhibition. Nature 606, 820–826. 10.1038/s41586-022-04814-x

Naritomi, Y., Fuchigami, S., 2013. Slow dynamics of a protein backbone in molecular dynamics simulation revealed by time-structure based independent component analysis. J. Chem. Phys. 139, 215102. 10.1063/1.4834695

Nayak, S.R., Joseph, D., Höfner, G., Dakua, A., Athreya, A., Wanner, K.T., Kanner, B.I., Penmatsa, A., 2023. Cryo-EM structure of GABA transporter 1 reveals substrate recognition and transport mechanism. Nat. Struct. Mol. Biol. 1–10. 10.1038/s41594-023-01011-w

Parrinello, M., Rahman, A., 1981. Polymorphic transitions in single crystals: A new molecular dynamics method. J. Appl. Phys. 52, 7182–7190. 10.1063/1.328693

Penmatsa, A., Wang, K.H., Gouaux, E., 2013. X-ray structure of dopamine transporter elucidates antidepressant mechanism. Nature 503, 85–90. 10.1038/nature12533

Radian, R., Kanner, B.I., 1983. Stoichiometry of sodium- and chloride-coupled .gamma.- aminobutyric acid transport by synaptic plasma membrane vesicles isolated from rat brain. Biochemistry 22, 1236–1241. 10.1021/bi00274a038

Scherer, M.K., Trendelkamp-Schroer, B., Paul, F., Pérez-Hernández, G., Hoffmann, M., Plattner, N., Wehmeyer, C., Prinz, J.-H., Noé, F., 2015. PyEMMA 2: A Software Package for Estimation, Validation, and Analysis of Markov Models. J. Chem. Theory Comput. 11, 5525–5542. 10.1021/acs.jctc.5b00743

Schicker, K., Uzelac, Z., Gesmonde, J., Bulling, S., Stockner, T., Freissmuth, M., Boehm, S., Rudnick, G., Sitte, H.H., Sandtner, W., 2012. Unifying concept of serotonin transporter-associated currents. J. Biol. Chem. 287, 438–445. 10.1074/jbc.M111.304261

Schousboe, A., Madsen, K.K., 2017. Delineation of the Role of Astroglial GABA Transporters in Seizure Control. Neurochem. Res. 42, 2019–2023. 10.1007/s11064-017-2188-x

Schousboe, A., Sarup, A., Larsson, O.M., White, H.S., 2004. GABA transporters as drug targets for modulation of GABAergic activity. Biochem. Pharmacol., Six Decades of GABA 68, 1557–1563. 10.1016/j.bcp.2004.06.041

Schrödinger, LLC, Warren DeLano, 2020. Pymol.

Singh, S.K., Piscitelli, C.L., Yamashita, A., Gouaux, E., 2008. A Competitive Inhibitor Traps LeuT in an Open-to-Out Conformation. Science 322, 1655–1661. 10.1126/science.1166777

Sohail, A., Jayaraman, K., Venkatesan, S., Gotfryd, K., Daerr, M., Gether, U., Loland, C.J., Wanner, K.T., Freissmuth, M., Sitte, H.H., Sandtner, W., Stockner, T., 2016. The Environment Shapes the Inner Vestibule of LeuT. PLOS Comput. Biol. 12, e1005197. 10.1371/journal.pcbi.1005197

Szöllősi, D., Stockner, T., 2022. Sodium Binding Stabilizes the Outward-Open State of SERT by Limiting Bundle Domain Motions. Cells 11, 255. 10.3390/cells11020255

Szöllősi, D., Stockner, T., 2021. Investigating the Mechanism of Sodium Binding to SERT Using Direct Simulations. Front. Cell. Neurosci. 15.

Tavoulari, S., Margheritis, E., Nagarajan, A., DeWitt, D.C., Zhang, Y.-W., Rosado, E., Ravera, S., Rhoades, E., Forrest, L.R., Rudnick, G., 2016. Two Na+ Sites Control Conformational Change in a Neurotransmitter Transporter Homolog*. J. Biol. Chem. 291, 1456–1471. 10.1074/jbc.M115.692012

van Meer, G., 1998. Lipids of the Golgi membrane. Trends Cell Biol. 8, 29–33. 10.1016/s0962-8924(97)01196-3

Varadi, M., Anyango, S., Deshpande, M., Nair, S., Natassia, C., Yordanova, G., Yuan, D., Stroe, O., Wood, G., Laydon, A., Žídek, A., Green, T., Tunyasuvunakool, K., Petersen, S., Jumper, J., Clancy, E., Green, R., Vora, A., Lutfi, M., Figurnov, M., Cowie, A., Hobbs, N., Kohli, P., Kleywegt, G., Birney, E., Hassabis, D., Velankar, S., 2022. AlphaFold Protein Structure Database: massively expanding the structural coverage of protein-sequence space with high-accuracy models. Nucleic Acids Res. 50, D439–D444. 10.1093/nar/gkab1061

Wassenaar, T.A., Ingólfsson, H.I., Böckmann, R.A., Tieleman, D.P., Marrink, S.J., 2015. Computational Lipidomics with insane: A Versatile Tool for Generating Custom Membranes for Molecular Simulations. J. Chem. Theory Comput. 11, 2144–2155. 10.1021/acs.jctc.5b00209

Wolf, M.G., Hoefling, M., Aponte-Santamaría, C., Grubmüller, H., Groenhof, G., 2010. g_membed: Efficient insertion of a membrane protein into an equilibrated lipid bilayer with minimal perturbation. J. Comput. Chem. 31, 2169–2174. 10.1002/jcc.21507

Yamashita, A., Singh, S.K., Kawate, T., Jin, Y., Gouaux, E., 2005. Crystal structure of a bacterial homologue of Na+/Cl--dependent neurotransmitter transporters. Nature 437, 215–223. 10.1038/nature03978

Yang, D., Gouaux, E., 2021. Illumination of serotonin transporter mechanism and role of the allosteric site. Sci. Adv. 7, eabl3857. 10.1126/sciadv.abl3857

Zhang, Y.-W., Tavoulari, S., Sinning, S., Aleksandrova, A.A., Forrest, L.R., Rudnick, G., 2018. Structural elements required for coupling ion and substrate transport in the neurotransmitter transporter homolog LeuT. Proc. Natl. Acad. Sci. 115, E8854–E8862. 10.1073/pnas.1716870115

Zhao, Y., Terry, D.S., Shi, L., Quick, M., Weinstein, H., Blanchard, S.C., Javitch, J.A., 2011. Substrate-modulated gating dynamics in a Na+-coupled neurotransmitter transporter homologue. Nature 474, 109–113. 10.1038/nature09971

Zhu, A., Huang, J., Kong, F., Tan, J., Lei, J., Yuan, Y., Yan, C., 2023. Molecular basis for substrate recognition and transport of human GABA transporter GAT1. Nat. Struct. Mol. Biol. 1–11. 10.1038/s41594-023-00983-z

Zomot, E., Bendahan, A., Quick, M., Zhao, Y., Javitch, J.A., Kanner, B.I., 2007. Mechanism of chloride interaction with neurotransmitter:sodium symporters. Nature 449, 726–730. 10.1038/nature06133

Zwanzig, R., 1983. From classical dynamics to continuous time random walks. J. Stat. Phys. 30, 255–262. 10.1007/BF01012300

