## Supplementary Information for "Interaction of GAT1 with sodium ions: from efficient recruitment to stabilisation of substrate and conformation"

**Affiliations:**
[1] Centre for Physiology and Pharmacology, Institute of Pharmacology, Medical University of Vienna, Waehrigerstr. 13A, 1090 Vienna, Austria
[2] Department of Theoretical and Computational Biophysics, Max Planck Institute for Multidisciplinary Sciences, Am Fassberg 11, 37077 Göttingen, Germany
[3] Department of Biochemistry and Molecular Biology, Institute for Medical Research Israel-Canada, Hebrew University, Hadassah Medical School, Jerusalem 91120, Israel
[4] Center for Addiction Research and Science-AddRess, Medical University Vienna, Waehringer Strasse 13A, 1090, Vienna, Austria
[5] Hourani Center for Applied Scientific Research, Al-Ahliyya Amman University, Amman, Jordan
[6] Institute of Pharmacology and Toxicology, University of Zurich, Zurich, Switzerland

**#corresponding author**
Thomas Stockner;

**$ Shared first authors**
These authors contributed evenly to the manuscript.

**Contents:**
Supplementary Figures, Molecular dynamics production parameters

**Supplementary Figure 1. Sequence alignment of SLC6 transporters:** The indicated sequence numbers are those of GAT1. The residue colour code is according to clustal. Fully conserved residues are indicated by a star, conserved residues by one or two dots. At the bottom the degree of conservation is indicated by a bar graph.

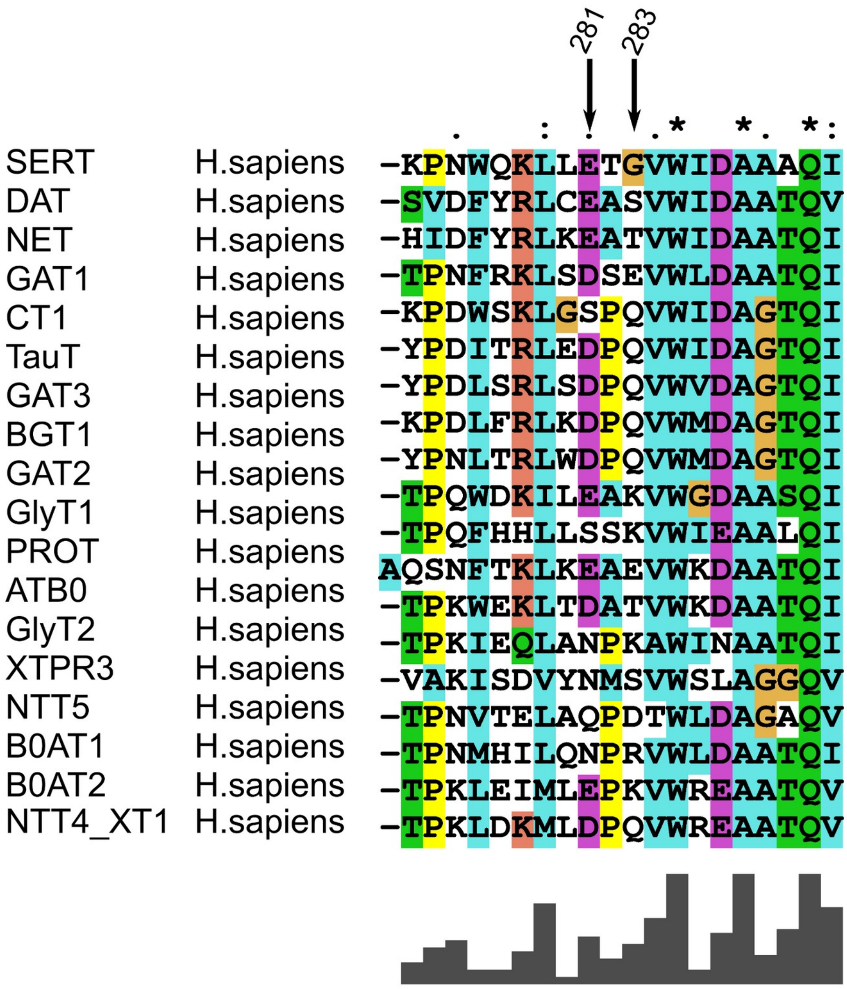

**Supplementary Figure 2a. Sodium Binding to NA1 of wild-type GAT1:** Time evolution of the distances of Na1 to the recruitment site, to the temporary site and to NA1. A rolling average of 5 nanoseconds was applied to smoothen the data.

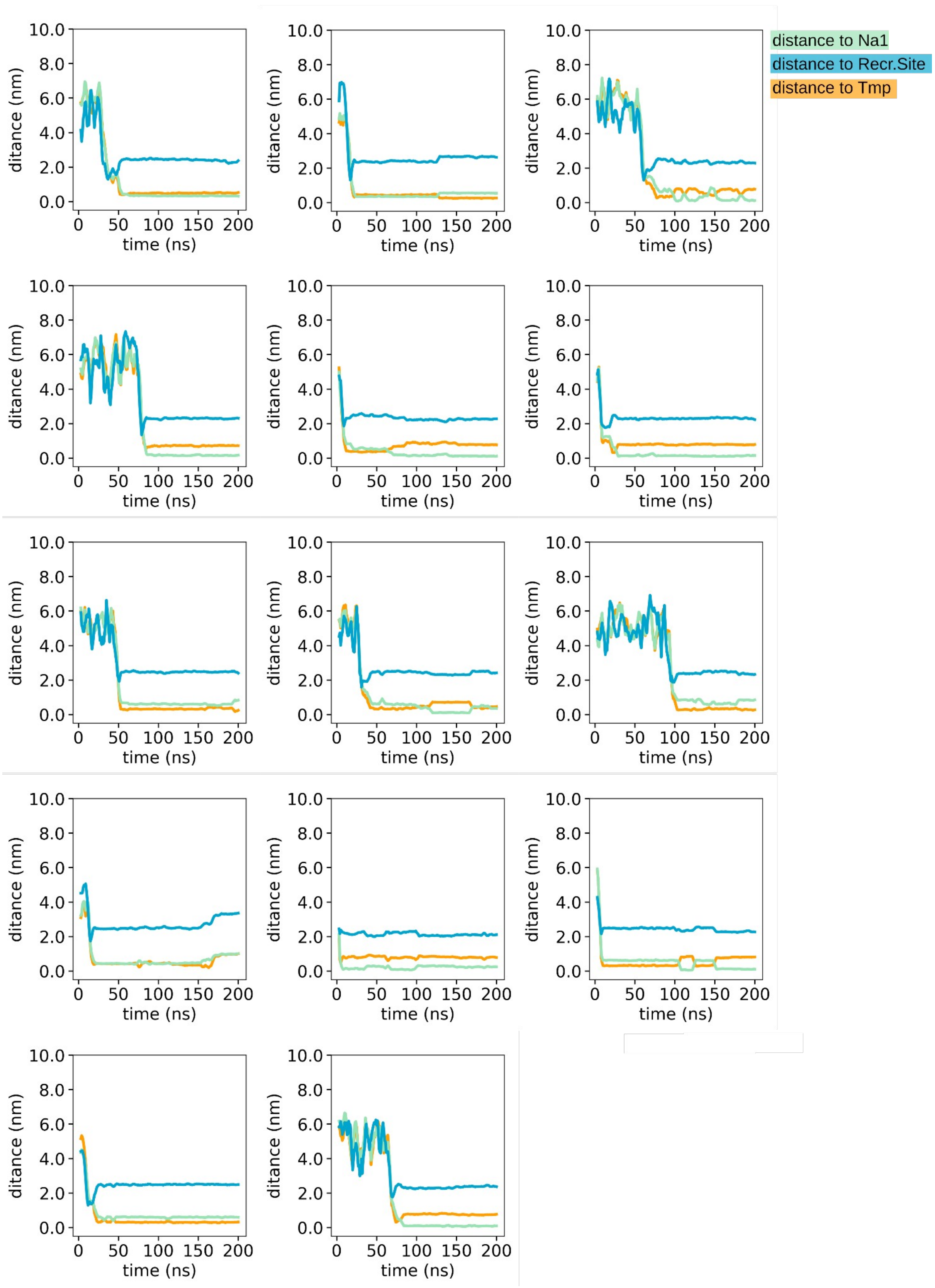

**Supplementary Figure 2b. Sodium Binding to NA2 of wild-type GAT1:** Time evolution of the distances of Na2 to the recruitment site, to the temporary site and to NA2. A rolling average of 5 nanoseconds was applied to smoothen the data.

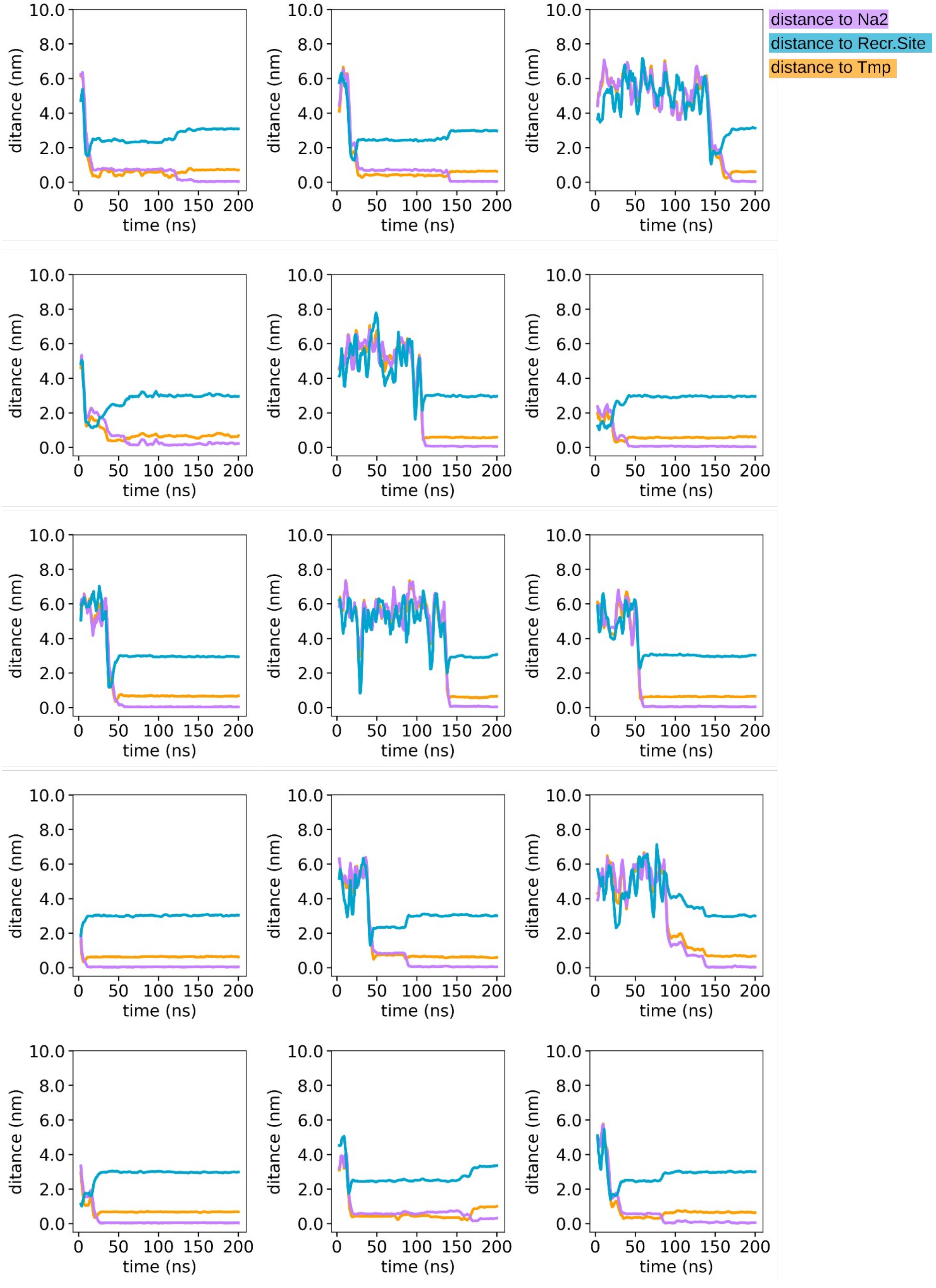

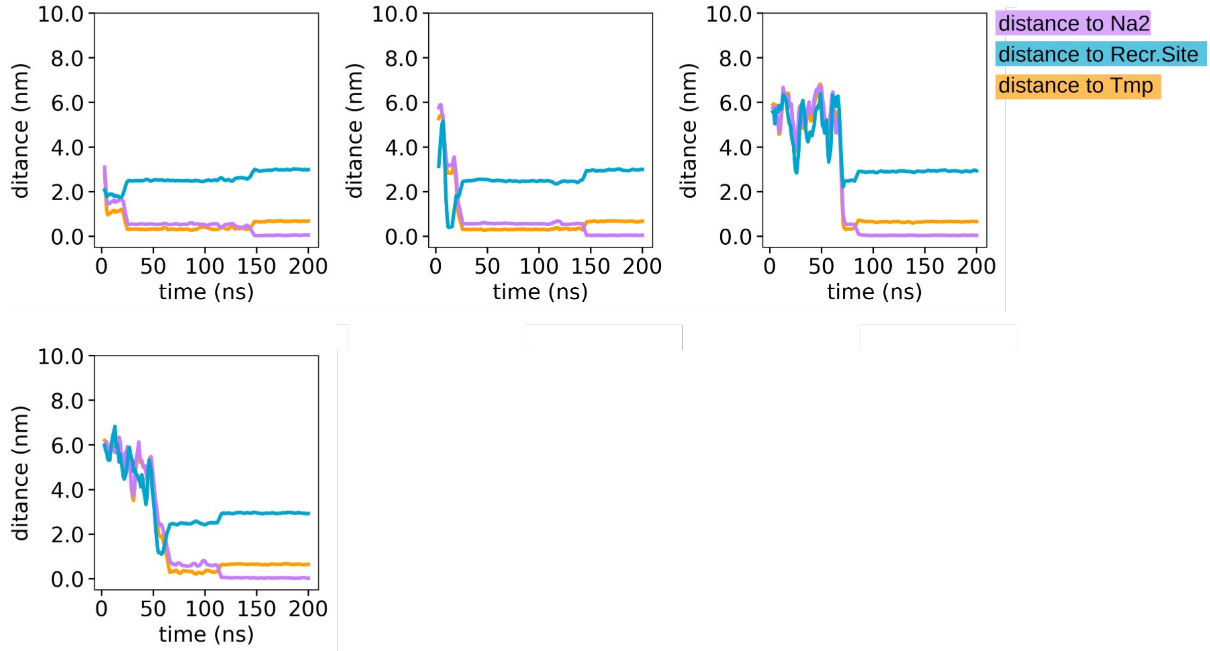
distance to Na2
distance to Recr.Site
distance to Tmp
ditance (nm)
time (ns)
10.0
8.0
6.0
4.0
2.0
0.0
0
50
100
150
200

**Supplementary Figure 2c. Sodium Binding to NA1 and NA2 of the double mutant (D281A-E283A) of GAT1:** Time evolution of the distances of Na1 and Na2 ions to the recruitment site, to the temporary site NA1 and to NA2. A rolling average of 5 nanoseconds was applied to smoothen the data.

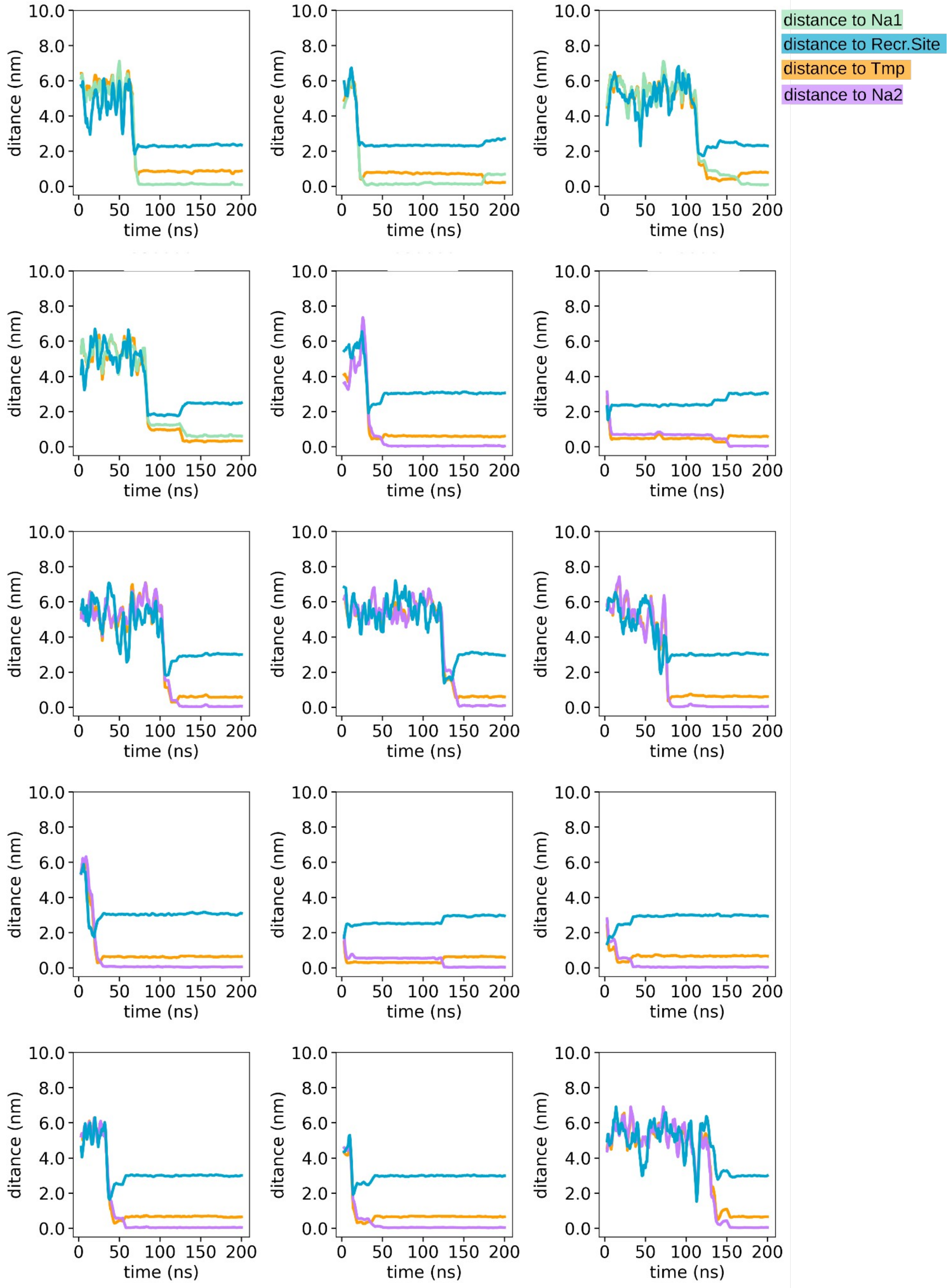

**Supplementary Figure 3. Compactness of the sodium binding sites.**

The compactness of the sodium binding sites NA1 and NA2 was assessed by determining the average distance from the centre of mass (CoM) of the sodium coordinating atoms to the respective sodium coordinating atoms. The time evolution shows the compactness of NA1 and NA2 per simulations, the vertical lines denote the time point in which sodium enters NA1 (cyan), NA2 (violet), or the temporary site (orange).

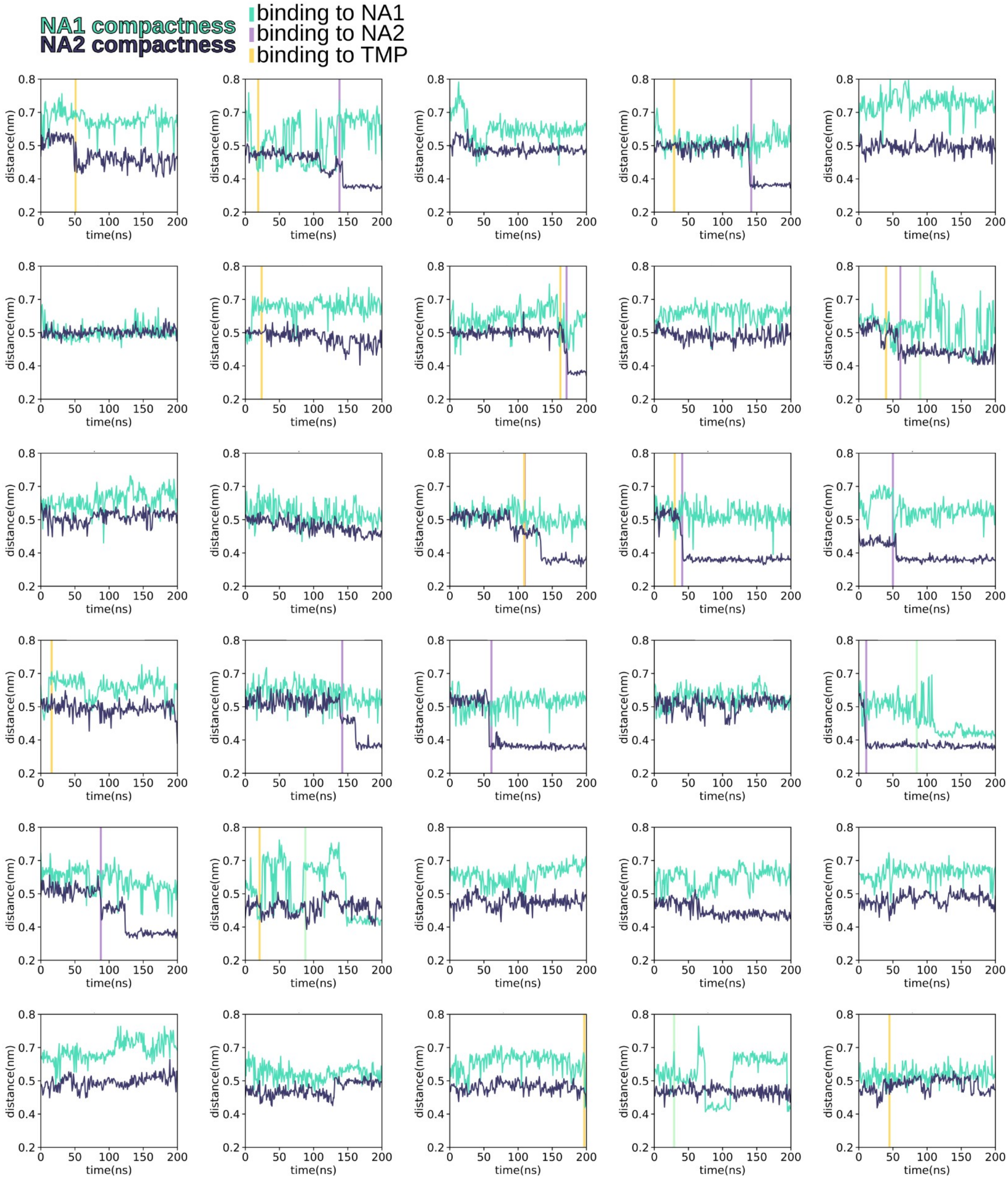

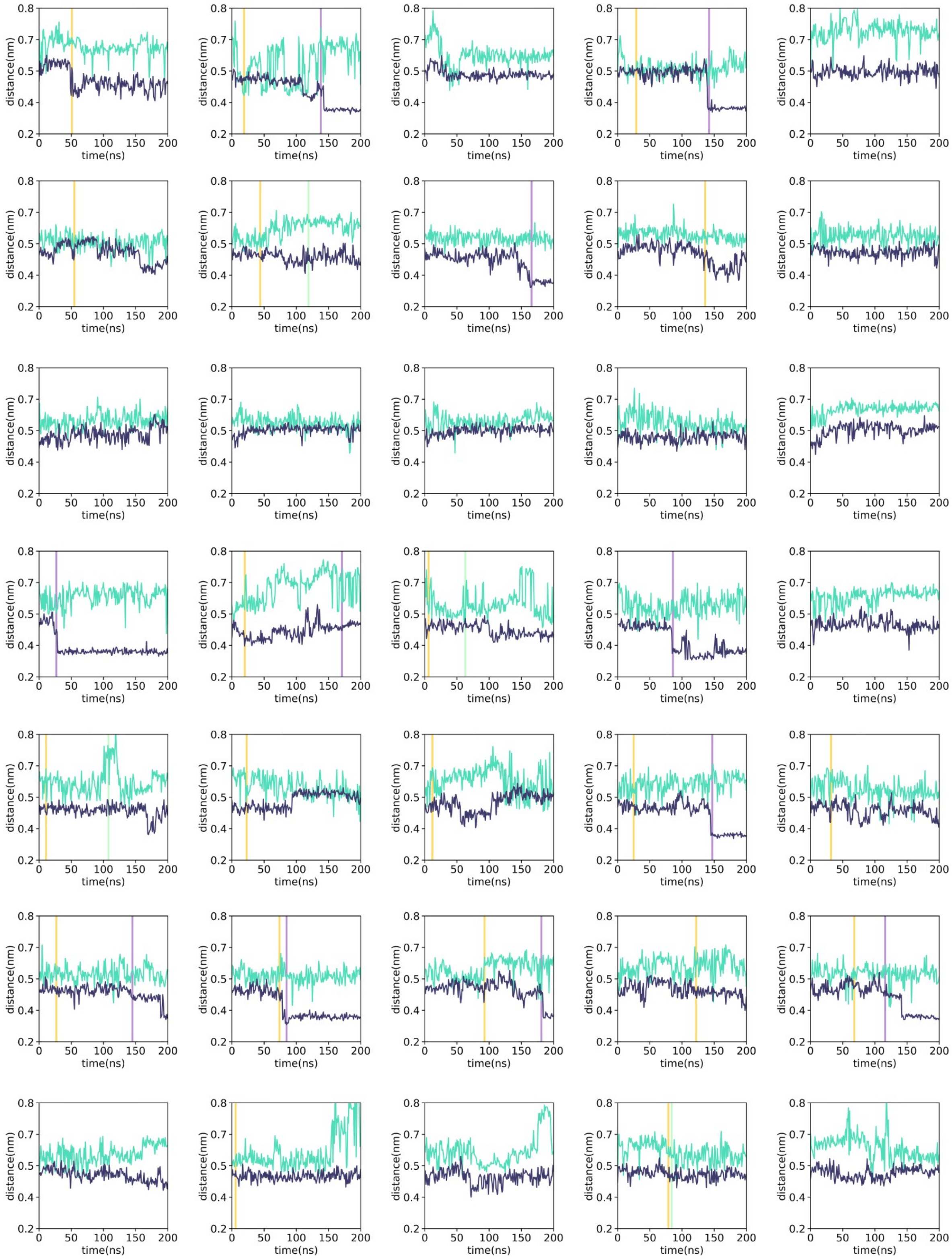
distance(nm)
time(ns)
0.2
0.4
0.5
0.7
0.8
0
50
100
150
200

**Supplementary Figure 4. Principal components analysis (PCA) of sodium binding:** The covariance matrix was calculated using all 60 trajectories of sodium binding to wild-type GAT1, using the C$\alpha$ atoms of the transmembrane helices (49:75, 64:76, 80:109, 121:155, 212:230, 282:309, 316:348,321345, 377:411, 420:439) as a fitting group. **a,b)** Subsequently, we divided the trajectories into two groups based on Na1 or Na2 binding and projected them onto the principal components previously calculated. The pink triangle corresponds to the NaCl-bound protein conformation (AF-based). **c)** The variance of the first fifty eigenvalues that describes the variance of each principal component**.**

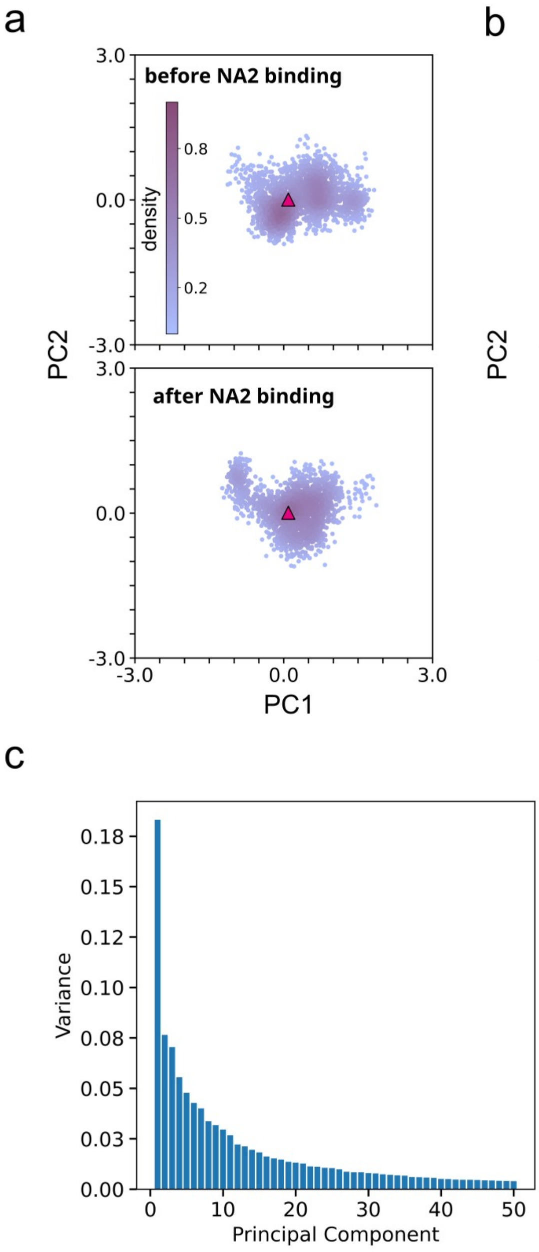

**Supplementary Figure 5. Markovian behaviour validation: a)** The Chapman-Kolmogorov (CK) test using a lag time equal to 25 ns and assuming four macrostates was used to evaluate Markovian behaviour. The predicted values obtained from the MSMs are shown by the solid lines while the estimated values for longer lag time are shown by the dashed traces. The superposition of predicted and estimated values indicates that the MSM assures Markovian behaviour. **b)** Time traces show the relation between the lag time and the implied timescales (or relaxation time) associated with the ten slowest processes, with the top blue trace indicating the slowest process. The solid lines refer to the maximum likelihood result and the dashed lines show the ensemble mean computed with Bayesian sampling procedure. The black line with the grey area indicates the timescale threshold where the MSM cannot resolve processes. In both panels, the non-grey areas indicate 95% confidence intervals computed using the Bayesian sampling procedure mentioned above.

a

b

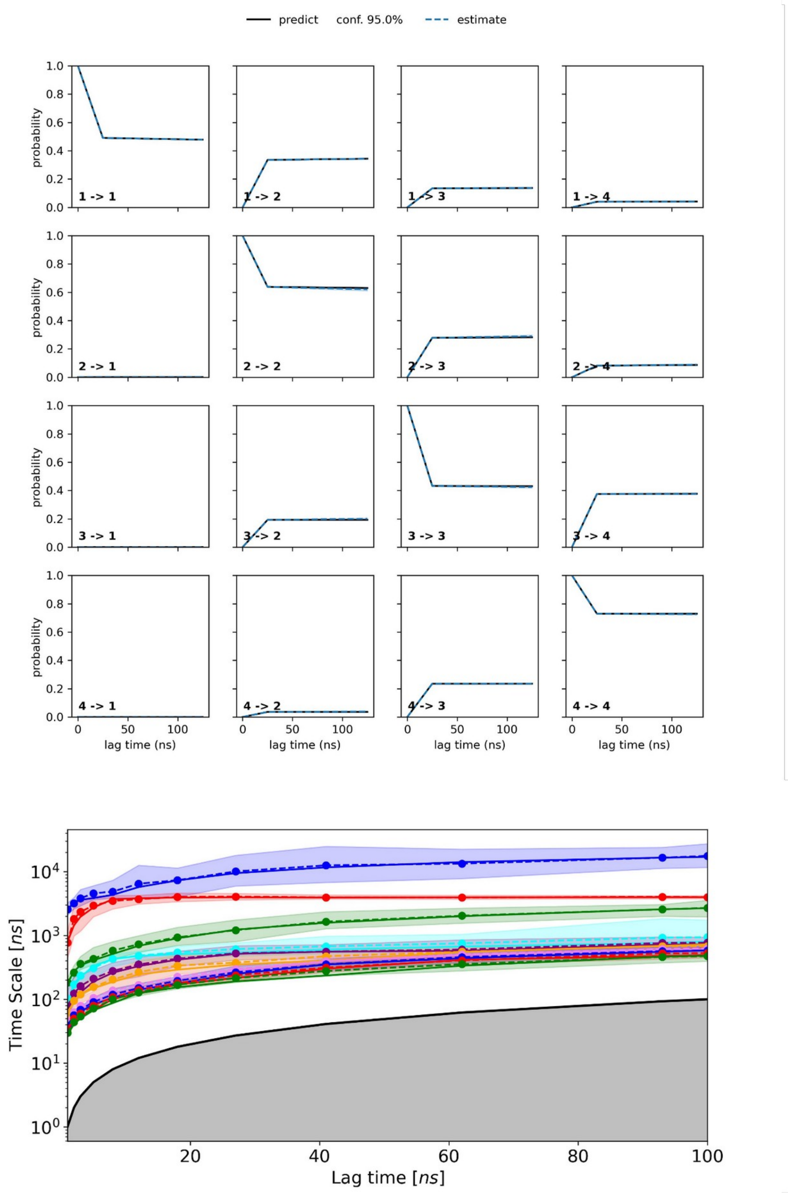

**MDP file:** Parameter file controlling the production simulation. The same parameters were applied to all simulations, only the total simulation length differed.

```
; VARIOUS PREPROCESSING OPTIONS
; Preprocessor information: use cpp syntax.
; e.g.: -I/home/joe/doe -I/home/mary/roe
;include                  =
; e.g.: -DPOSRES -DFLEXIBLE (note these variable names are case sensitive)
;define                   = -DPOSRES

; RUN CONTROL PARAMETERS
integrator               = md
; Start time and timestep in ps
tinit                    = 0
dt                       = 0.002
nsteps                   = 10000000 ;200 ns
; For exact run continuation or redoing part of a run
init_step                = 0
; Part index is updated automatically on checkpointing (keeps files separate)
simulation_part          = 1
; mode for center of mass motion removal
comm-mode                = linear
; number of steps for center of mass motion removal
nstcomm                  = 100
; group(s) for center of mass motion removal
comm-grps                = ProtLigIon membrane Water_and_ions

; LANGEVIN DYNAMICS OPTIONS
; Friction coefficient (amu/ps) and random seed
bd-fric                  = 0
ld-seed                  = 1993

; ENERGY MINIMIZATION OPTIONS
; Force tolerance and initial step-size
emtol                    = 1000
emstep                   = 0.0001
; Max number of iterations in relax-shells
niter                    = 20
; Step size (ps^2) for minimization of flexible constraints
fcstep                   = 0.001
; Frequency of steepest descents steps when doing CG
nstcgsteep               = 50
nbfgscorr                = 100
```

```
; TEST PARTICLE INSERTION OPTIONS
rtpi                     = 0.05

; OUTPUT CONTROL OPTIONS
; Output frequency for coords (x), velocities (v) and forces (f)
nstxout                  = 5000 ; 10 ps
nstvout                  =
nstfout                  = 0
; Output frequency for energies to log file and energy file
nstlog                   = 1000
nstcalcenergy            = 100
nstenergy                = 1000
; Output frequency and precision for .xtc file
nstxout-compressed       = 0
compressed-x-precision   = 1000
; This selects the subset of atoms for the compressed
; trajectory file. You can select multiple groups. By
; default, all atoms will be written.
compressed-x-grps        =
; Selection of energy groups
energygrps               =

; NEIGHBORSEARCHING PARAMETERS
; cut-off scheme (Verlet: particle based cut-offs, group: using
charge groups)
cutoff-scheme            = Verlet
; nblist update frequency
nstlist                  = 50
; ns algorithm (simple or grid)
ns-type                  = Grid
; Periodic boundary conditions: xyz, no, xy
pbc                      = xyz
periodic_molecules       = no
; Allowed energy error due to the Verlet buffer in kJ/mol/ps per
atom,
; a value of -1 means: use rlist
verlet-buffer-tolerance  = 0.005
; nblist cut-off
rlist                    = 0.9
; long-range cut-off for switched potentials
rlistlong                = -1
nstcalclr                = -1

; OPTIONS FOR ELECTROSTATICS AND VDW
; Method for doing electrostatics
```

```
coulombtype              = PME
coulomb-modifier         = Potential-shift-Verlet
rcoulomb-switch          =
rcoulomb                 = 0.9
; Relative dielectric constant for the medium and the reaction field
epsilon_r                = 1.0
epsilon_rf               = 1
; Method for doing Van der Waals
vdw-type                 = Cut-off
vdw-modifier             = Potential-shift-Verlet
; cut-off lengths
rvdw-switch              =
rvdw                     = 0.9
; Apply long range dispersion corrections for Energy and Pressure
DispCorr                 = EnerPres
; Extension of the potential lookup tables beyond the cut-off
table-extension          = 1
; Separate tables between energy group pairs
energygrp-table          =
; Spacing for the PME/PPPM FFT grid
fourierspacing           = 0.12
; FFT grid size, when a value is 0 fourierspacing will be used
fourier_nx               = 0
fourier_ny               = 0
fourier_nz               = 0
; EWALD/PME/PPPM parameters
pme_order                = 4
ewald_rtol               = 1e-05
ewald-rtol-lj            = 0.001
lj-pme-comb-rule         = Geometric
ewald_geometry           = 3d
epsilon_surface          = 0

; IMPLICIT SOLVENT ALGORITHM
implicit_solvent         = No

; GENERALIZED BORN ELECTROSTATICS
; Algorithm for calculating Born radii
gb-algorithm             = Still
; Frequency of calculating the Born radii inside rlist
nstgbradii               = 1
; Cutoff for Born radii calculation; the contribution from atoms
; between rlist and rgbradii is updated every nstlist steps
rgbradii                 = 1
; Dielectric coefficient of the implicit solvent
gb-epsilon-solvent       = 80
```

```
; Salt concentration in M for Generalized Born models
gb-saltconc              = 0
; Scaling factors used in the OBC GB model. Default values are
OBC(II)
gb-obc-alpha             = 1
gb-obc-beta              = 0.8
gb-obc-gamma             = 4.85
gb-dielectric-offset     = 0.009
sa-algorithm             = Ace-approximation
; Surface tension (kJ/mol/nm^2) for the SA (nonpolar surface) part
of GBSA
; The value -1 will set default value for Still/HCT/OBC GB-models.
sa-surface-tension       = -1

; OPTIONS FOR WEAK COUPLING ALGORITHMS
; Temperature coupling
tcoupl                   = v-rescale
nsttcouple               = -1
nh-chain-length          = 10
print-nose-hoover-chain-variables = no
; Groups to couple separately
tc-grps                  = ProtLigIon membrane Water_and_ions
; Time constant (ps) and reference temperature (K)
tau-t                    = 0.5   0.5   0.5
ref-t                    = 310   310   310
; pressure coupling
Pcoupl                    = Parrinello-Rahman ; Berendsen for EM and
equilibration;Parrinello-Rahman for production
Pcoupltype               = Semiisotropic
nstpcouple               = -1
; Time constant (ps), compressibility (1/bar) and reference P (bar)
tau-p                     = 20.1       ; Use 5 for Berendsen; 20 for
Parrinello-Rahman;
compressibility          = 4.5e-05   4.5e-05
ref-p                    = 1.0       1.0
; Scaling of reference coordinates, No, All or COM
refcoord_scaling         = All


; SIMULATED ANNEALING
; Type of annealing for each temperature group (no/single/periodic)
annealing                = no
; Number of time points to use for specifying annealing in each
group
annealing-npoints        =
; List of times at the annealing points for each group
```

```
annealing-time           =
; Temp. at each annealing point, for each group.
annealing-temp           =

; GENERATE VELOCITIES FOR STARTUP RUN
gen-vel                  = no
gen-temp                 = 310.0
gen-seed                 = -1

; OPTIONS FOR BONDS
constraints              = h-bonds
; Type of constraint algorithm
constraint-algorithm     = lincs
; Do not constrain the start configuration
continuation             = no
; Use successive overrelaxation to reduce the number of shake
iterations
Shake-SOR                = yes
; Relative tolerance of shake
shake-tol                = 0.0001
; Highest order in the expansion of the constraint coupling matrix
lincs-order              = 4
; Number of iterations in the final step of LINCS. 1 is fine for
; normal simulations, but use 2 to conserve energy in NVE runs.
; For energy minimization with constraints it should be 4 to 8.
lincs-iter               = 2
; Lincs will write a warning to the stderr if in one step a bond
; rotates over more degrees than
lincs-warnangle          = 30
; Convert harmonic bonds to morse potentials
morse                    = no

; ENERGY GROUP EXCLUSIONS
; Pairs of energy groups for which all non-bonded interactions are
excluded
energygrp-excl           =

; WALLS
; Number of walls, type, atom types, densities and box-z scale
factor for Ewald
nwall                    = 0
wall_type                = 9-3
wall_r_linpot            = -1
wall-atomtype            =
wall-density             =
wall_ewald_zfac          = 3
```

```
; COM PULLING
; Pull type: no, umbrella, constraint or constant-force
pull                     = no

; ENFORCED ROTATION
; Enforced rotation: No or Yes
rotation                 = no

; Group to display and/or manipulate in interactive MD session
IMD-group                =

; NMR refinement stuff
; Distance restraints type: No, Simple or Ensemble
disre                    = No
; Force weighting of pairs in one distance restraint: Conservative
or Equal
disre-weighting          = Conservative
; Use sqrt of the time averaged times the instantaneous violation
disre-mixed              = no
disre-fc                 = 100
disre-tau                = 0
; Output frequency for pair distances to energy file
nstdisreout              = 5000
; Orientation restraints: No or Yes
orire                    = no
; Orientation restraints force constant and tau for time averaging
orire-fc                 = 0
orire-tau                = 0
orire-fitgrp             =
; Output frequency for trace(SD) and S to energy file
nstorireout              = 100

; Free energy variables
free-energy              = no
couple-moltype           =
couple-lambda0           = vdw-q
couple-lambda1           = vdw-q
couple-intramol          = no
init-lambda              = 0
init-lambda-state        = -1
delta-lambda             = 0
nstdhdl                  = 50
fep-lambdas              =
mass-lambdas             =
coul-lambdas             =
```

```
vdw-lambdas              =
bonded-lambdas           =
restraint-lambdas        =
temperature-lambdas      =
calc-lambda-neighbors    = 1
init-lambda-weights      =
dhdl-print-energy        = no
sc-alpha                 = 0
sc-power                 = 1
sc-r-power               = 6
sc-sigma                 = 0.3
sc-coul                  = no
separate-dhdl-file       = yes
dhdl-derivatives         = yes
dh_hist_size             = 0
dh_hist_spacing          = 0.1

; Non-equilibrium MD stuff
acc-grps                 =
accelerate               =
freezegrps               =
freezedim                =
cos-acceleration         = 0
deform                   =

; simulated tempering variables
simulated-tempering      = no
simulated-tempering-scaling = geometric
sim-temp-low             = 300
sim-temp-high            = 300

; Electric fields
; Format is number of terms (int) and for all terms an amplitude
(real)
; and a phase angle (real)
E-x                      =
; Time dependent (pulsed) electric field. Format is omega, time for
pulse
; peak, and sigma (width) for pulse. Sigma = 0 removes pulse,
leaving
; the field to be a cosine function.
E-xt                     =
E-y                      =
E-yt                     =
E-z                      =
E-zt                     =
```

```
; Ion/water position swapping for computational electrophysiology
setups
; Swap positions along direction: no, X, Y, Z
swapcoords               = no

; AdResS parameters
adress                   = no

; User defined thingies
user1-grps               =
user2-grps               =
userint1                 = 0
userint2                 = 0
userint3                 = 0
userint4                 = 0
userreal1                = 0
userreal2                = 0
userreal3                = 0
userreal4                = 0
```
